# Disentangling neuronal inhibition and inhibitory pathways in the lateral habenula

**DOI:** 10.1101/633271

**Authors:** Jack F. Webster, Rozan Vroman, Kira Balueva, Peer Wulff, Shuzo Sakata, Christian Wozny

## Abstract

The lateral habenula (LHb) is hyperactive in depression, and thus potentiating inhibition of this structure makes an interesting target for future antidepressant therapies. However, the circuit mechanisms mediating inhibitory signalling within the LHb are not well-known. We addressed this issue by studying LHb neurons expressing either parvalbumin (PV), neuron-derived neurotrophic factor (Ndnf) or somatostatin (SOM), three markers of particular sub-classes of neocortical inhibitory neurons. While we report that Ndnf is not representative of any particular sub-population of LHb neuron, we find that both PV and SOM are expressed by physiologically distinct sub-classes. Furthermore, we describe multiple sources of inhibitory input to the LHb arising from both local PV-positive neurons, and from PV-positive neurons in the medial dorsal thalamic nucleus, and from SOM-positive neurons in the ventral pallidum. These findings hence provide new insight into inhibitory control within the LHb, and highlight that this structure is more neuronally diverse than previously thought.

**Summary:** The lateral habenula receives inhibitory input from three distinct sources: from local PV-positive neurons, from PV-positive neurons in the medial dorsal thalamic nucleus (MDT); and from SOM-positive neurons in the ventral pallidum (VP).

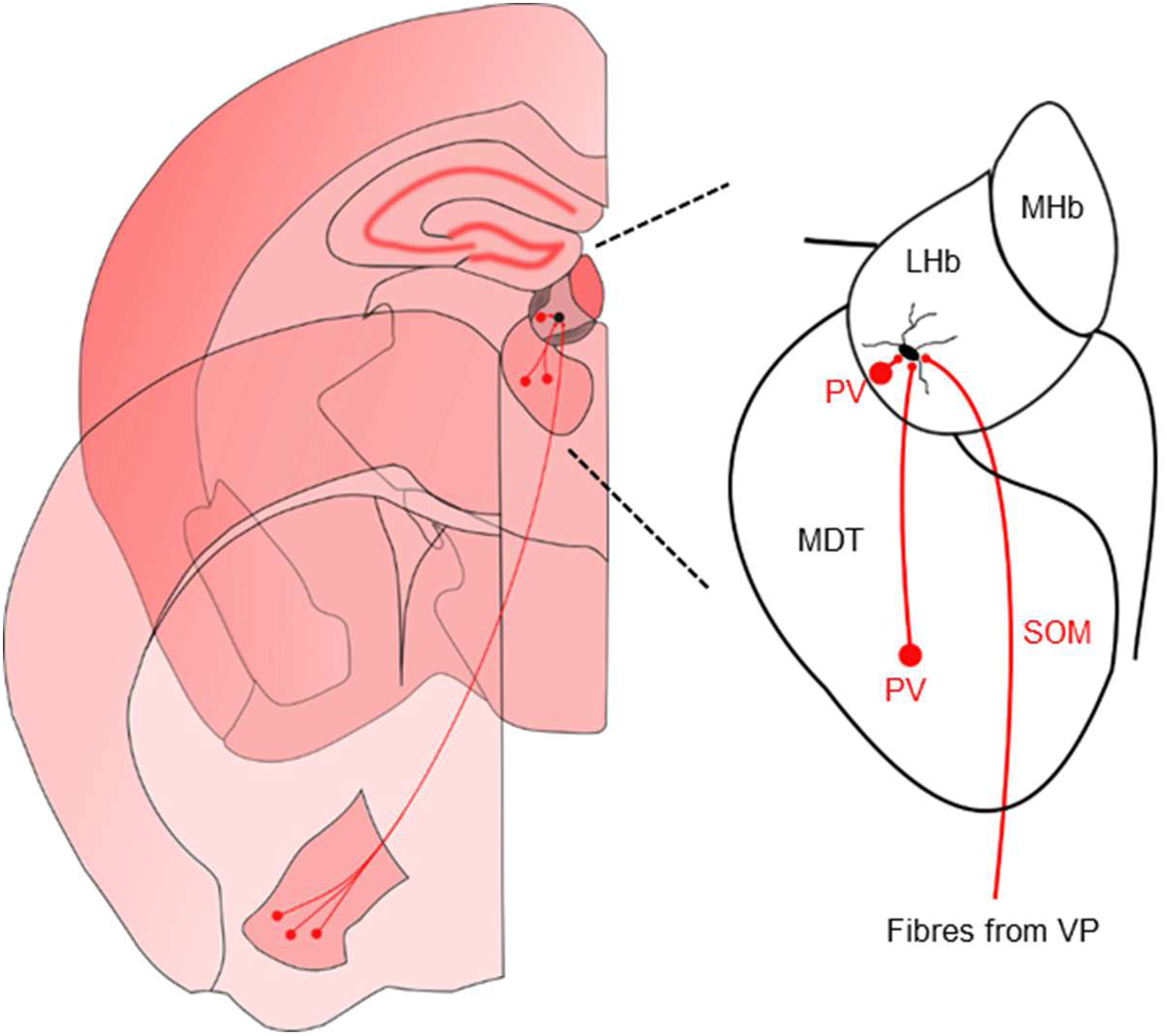

**Significance statement:** The circuitry by which inhibitory signalling is processed within the lateral habenula is currently not well understood; yet this is an important topic as inhibition of the lateral habenula has been shown to have antidepressant efficacy. We therefore investigated inhibitory signalling mechanisms within the lateral habenula by studying input neurons expressing markers commonly associated with inhibitory identity. We identity sources of inhibitory input from both local neurons, and arising from neurons in the medial dorsal thalamic nucleus and ventral pallidum.

**Contributions:** J.F.W. performed the experiments. R.V. contributed to experiments. J.F.W. analysed the data. K.B. and P.W. designed and performed the in situ hybridisation experiments. C.W. designed and supervised the study, and helped J.F.W write the manuscript. R.V. and S.S. contributed to the manuscript and discussions.

## Introduction

The lateral habenula (LHb) is an epithalamic brain structure, which acts as an inhibitory modulator of the midbrain reward circuitry, including the ventral tegmental area (VTA) (Christoph et al., 1986; Ji and Shepard, 2007; Matsumoto and Hikosaka, 2007) and the raphe nuclei (Wang and Aghajanian, 1977). Recently, the LHb has drawn renewed attention (Geisler and Trimble, 2008) due to the revelation that it becomes pathologically hyperactive in major depressive disorder (MDD) (Cui et al., 2018; Lecca et al., 2016; Li et al., 2011; Sartorius et al., 2010; Shabel et al., 2014; Tchenio et al., 2017; Yang et al., 2018), thus providing excessive inhibition to these reward centres and silencing the associated positive emotions. Concurrently, this makes the LHb an intriguing target for novel antidepressant therapies (Cui et al., 2018; Lecca et al., 2016; Li et al., 2011; Sartorius et al., 2010; Schneider et al., 2013; Winter et al., 2011; Yang et al., 2018). Notably, multiple studies have shown that potentiation of inhibition within the LHb yields an antidepressant effect (Huang et al., 2019; Lecca et al., 2016; Tchenio et al., 2017; Winter et al., 2011), likely by disinhibition of the reward circuitry. Despite this, however, the neural circuitry which mediates inhibitory signalling within the LHb is not well understood. To date, very few populations of local inhibitory neurons have been identified within the LHb (Wagner et al., 2016a; Weiss and Veh, 2011; Zhang et al., 2018), and external inhibitory input appears to arise primarily from GABA / glutamate co-releasing fibres (Meye et al., 2016; Root et al., 2014; Shabel et al., 2014; Wallace et al., 2017). Yet the LHb does receive a large inhibitory input (Lecca et al., 2016; Wagner et al., 2016a), and recently some exclusively GABAergic fibres have been reported to project to it (Barker et al., 2017; Faget et al., 2018; Huang et al., 2019). It is hence likely that other as-of-yet undefined circuitry components mediate inhibitory signalling within the LHb, and studying these may yield further insights into the underlying mechanisms by which it is implicated in the pathogenesis of MDD.

We therefore sought to address this issue of inhibitory control within the LHb. Both parvalbumin (PV) and somatosatin (SOM) have long been accepted as a neuronal marker associated with particular sub-populations of inhibitory interneurons within many structures throughout the brain including the neocortex (Tremblay et al., 2016), hippocampus (Klausberger and Somogyi, 2008) and striatum (Tepper et al., 2011). Neuron-derived neurotrophic factor (Ndnf) has recently been proposed to act as a selective marker for neocortical neurogliaform cells (Abs et al., 2018; Tasic et al., 2018, 2016), which are thought to be present within the LHb (Wagner et al., 2016a; Weiss and Veh, 2011), while neuropeptide Y (NPY) is also an accepted, although not entirely selective, marker of these neurons (Overstreet-Wadiche and McBain, 2015).

Referring to the question of inhibitory control within the LHb, we therefore asked if these markers also represented distinct sub-populations of inhibitory neurons within this structure, and aimed to characterise the circuitry formed by neurons expressing them. In contrast to other brain regions, we find that Ndnf and NPY are not confined to any particular sub-population of LHb neuron. Strikingly, however, we report three sources of inhibitory input to the LHb arising from locally-targeting PV-positive neurons within the LHb, from PV-positive neurons within the medial dorsal thalamic nucleus, and from SOM-positive neurons in the ventral pallidum.

## Materials and Methods

### Animals

All procedures were approved by the Ethics committee of the University of Strathclyde, Glasgow in accordance with the relevant UK legislation (the Animals (Scientific Procedures) Act, 1986). Male and female mice from each strain were used in this work. All animals were maintained on a C57BL/6 background, and kept on a 12:12 light/dark cycle under standard group housing conditions with unlimited access to water and normal mouse chow. New-born pups were housed with parents until weaning at P21. To generate transgenic reporter-bearing offspring, transgenic mice of the Ndnf-IRES-Cre (Jax. ID 025836) (Tasic et al., 2016), PV-IRES-Cre (Jax. ID 017320) (Hippenmeyer et al., 2005) or SOM-IRES-Cre (Jax. ID 018973) (Taniguchi et al., 2011) driver lines were crossed with either Ai32 (Jax. ID 025109) (Madisen et al., 2012) or Ai9 (Jax. ID 007909) (Madisen et al., 2010) reporter mice driving expression of Channelrhodospin-2 (ChR2) and enhanced yellow fluorescent protein (eYFP), or the enhanced red fluorescent protein variant tdTomato in a Cre-dependent manner, respectively. The resulting offspring strains are hence referred to as: Ndnf-IRES-Cre::Ai32, PV-IRES-Cre::Ai32, SOM-IRES-Cre::Ai32, Ndnf-IRES-Cre::Ai9, PV-IRES-Cre::Ai9 and SOM-IRES-Cre::Ai9. NPY-hrGFP mice (van den Pol et al., 2009) were also used in this study.

### Production of recombinant AAV vectors

pAAV-Ef1a-DIO hChR2(E123T/T159C)-EYFP plasmid (a gift from Karl Deisseroth; RRID:Addgene_35509) was packaged into AAVs as described previously (Murray et al., 2011). Briefly, virions containing a 1:1 ratio type 1 and type 2 capsid proteins were produced by transfecting human embryonic kidney (HEK) 293 cells with the rAAV backbone plasmid pAAV-Ef1a-DIO hChR2(E123T/T159C)-EYFP along with AAV1 (pH21), AAV2 (pRV1) and adenovirus helper plasmid pFdelta6 using the calcium phosphate method. 48 hours post transfection, cells were harvested and rAAVs were purified using 1 mL HiTrap heparin columns (GE Healthcare Bio-Sciences, Uppsala, SwedenSigma) and concentrated using Amicon Ultra centrifugal filter devices (Merck Millipore, Tullagreen, Ireland). Infectious rAAV particles (viral titer) were calculated by serially infecting HEK293 cells stably expressing Cre-recombinase and counting GFP722 positive cells.

### Stereotaxic viral injections

PV-IRES-Cre mice (P31-48) or SOM-IRES-Cre mice (P34-44) were deeply anaesthetized via inhaled isoflurane (5 % for induction; 1-2 % for maintenance), transferred to a stereotaxic frame (Narishige, Tokyo, Japan) and were subcutaneously injected with the analgesics carprofen (5 mg/kg) in the nape and lidocaine (4 mg/kg) under the scalp. Intracranial injections were made using a glass micropipette pulled using a PC-100 vertical puller (Narishige, Tokyo, Japan). Under aseptic conditions, the skull was exposed and a small burr hole was drilled above the habenula within the left hemisphere. The injection capillary was then advanced, and a Cre-dependent viral vector was injected into the LHb, the medial dorsal thalamic nucleus (MDT) or the ventral pallidum (VP) at a rate of 25 nL / min using a pressure microinjector (Narishige, Tokyo, Japan). Two different vector solutions were used in this study: one containing a pAAV-EF1α-Switch:NLSmRuby2/ChR2(H134R)-EYFP-HGHpA (Wozny et al., 2018) (titre: 2 x 10^13^ infectious particles / mL), an adeno-associated virus containing capsid protein 9 which drives expression of ChR2 and eYFP in Cre-expressing cells, or the red fluorescent protein variant mRuby2 in the absence of Cre; or AAV Ef1α-DIO-hChR2-eYFP (titre: 1.6 x 10^8^ infectious particles / mL), a mixture of adeno-associated virus particles of serotypes 1 and 2 containing a 1:1 ratio of capsid proteins type 1 and 2. Parameters for LHb-targeted injections were: coordinates (from Bregma, in mm) AP −1.5, ML −0.4, depth 3; volume 50-100 nL; N = 8 PV-IRES-Cre mice, N = 4 SOM-IRES-Cre mice. Parameters for MDT-targeted injections were: AP −1.355, ML −0.75 depth 3.5; volume 200 nL; N = 2 mice or AP −1.155, ML −0.45 depth 3.7; volume 200 nL; N = 4 mice. Note that for the MDT-targeted injection coordinates were adjusted following data acquisition from the first round of experiments. Parameters for VP-targeted injections were: AP 0.145, ML −1.55, depth 5.65; volume 100-250 nL; N = 3 mice. Following injection, the needle was left for 10 minutes to allow the virus to diffuse before being slowly withdrawn. Animals were allowed to recover from anaesthesia on a heat pad. Following completion of surgery, animals were given at least two weeks to allow expression of the virus before acute slice preparation for electrophysiology. Upon completion of electrophysiology, viral spread was assessed by imaging on a Leica SP5 or SP8 confocal microscope.

### Acute brain slice preparation

C57BL/6 (N = 5; P21-28), Ndnf-IRES-Cre::Ai32 (N = 6; P21-28), Ndnf-IRES-Cre::Ai9 (N = 5; P21-36), PV-IRES-Cre::Ai32 (N = 19; P23-40), PV-IRES-Cre::Ai9 (N = 8; P23-33), SOM-IRES-Cre::Ai32 (N = 5; P19-37), SOM-IRES-Cre::Ai9 (N = 3; P24-28) or surgically injected PV-IRES-Cre or SOM-IRES-Cre mice (described above) were humanely euthanized by cervical dislocation and immediately decapitated, and brains were rapidly removed and transferred to ice-cold oxygenated (95% O_2_; 5% CO_2_) sucrose-based artificial cerebro-spinal fluid (ACSF) solution containing (in mM): sucrose 50, NaCl 87, NaHCO_3_ 25, KCl 3, NaH_2_PO_4_ 1.25, CaCl_2_ 0.5, MgCl_2_ 3, sodium pyruvate 3 and glucose 10. Brains sections containing the habenula were then cut in the coronal plane at 250-300 µm on a Leica VT1200S vibratome (Leica Biosystems, Newcastle-upon-Tyne, UK). In order to ensure slices contained the habenula, the hippocampus was used as a visual guidance due to the easily identifiable structure and immediate proximity to the habenula. Following sectioning, slices were incubated in oxygenated sucrose-based ACSF at 35 °C for 30 minutes, and then incubated for a further 30 minutes at room temperature in ACSF containing (in mM) NaCl 115, NaHCO_3_ 25, KCl 3, NaH_2_PO_4_ 1.25, CaCl_2_ 2, MgCl_2_ 1, sodium pyruvate 3 and glucose 10. Following the incubation period, slices were stored at room temperature in oxygenated ACSF.

### Electrophysiological recordings

Individual slices were transferred to a recording chamber and continually perfused with oxygenated ACSF at a flow rate of 2-3 mL / min, and visualized with a Luigs and Neumann LN-Scope System (Luigs and Neumann, Ratingen, Germany). The habenula is easily identifiable under differential interference contrast microscopy even at low magnification and hence a 4X objective was used to locate the lateral habenular nucleus. A 60X objective was then used to identify suitable cells for whole-cell recordings. In the case of transgenic Ndnf-IRES-Cre::Ai9, PV-IRES-Cre::Ai9 or SOM-IRES-Cre::Ai9 slices, TdTomato-expressing cells could be selectively visualized with an Olympus XM10 fluorescent camera (Olympus, Southend-on-Sea, UK) upon photostimulation with a blue LED (pE-300^ultra^, Cool LED, Andover, UK). Recordings were made with a Multiclamp 700B Amplifier (Molecular Devices, California, USA). Glass micropipettes were filled with a solution containing (in mM) potassium gluconate 125, HEPES 10, KCl 6, EGTA 0.2, MgCl_2_ 2, Na-ATP 2, Na-GTP 0.5, sodium phosphocreatine 5, and with 0.2 % biocytin. pH was adjusted to 7.2 with KOH. For spontaneous current measurement experiments, a reduced chloride intracellular solution was used consisting of (in mM) potassium gluconate 140, potassium chloride 2, EGTA 0.2, Hepes 10, NaATP 2, NaGTP 0.5 and sodium phosphocreatine 5.

Once in whole-cell patch mode, the intrinsic properties of LHb neurons were assessed in current-clamp configuration using a stepping protocol consisting of 1 s long injections of increasing current (range: −250-250 pA; step size: 5-50 pA for LHb neurons and −500-1000 pA; step size 100 pA for cortical neurons). Action potential firing pattern was assessed in response to depolarizing current injection, while hyperpolarizing current injection allowed the characterisation of rebound action potential firing of neurons. Resting membrane potential (RMP) was assessed by recording the spontaneous activity of each neuron with no current injection for at least 30 seconds, while membrane input resistance was monitored by injecting a small hyperpolarizing pulse (100 ms; −10 to −100 pA) and measuring the voltage change. Spontaneous currents were observed in voltage clamp at a holding potential of −60 mV. Series resistance was monitored throughout. All neuronal voltage and current signals were low pass-filtered at 2-10 kHz and acquired at 10-25 kHz using an ITC-18 digitizer interface (HEKA, Pfalz, Germany). The data acquisition software used was Axograph X.

### Optogenetic experiments and pharmacology

For optogenetic experiments, acute brain slices were prepared from transgenic Ndnf-IRES-Cre::Ai32, PV-IRES-Cre::Ai32 and SOM-IRES-Cre::Ai32 offspring as above in darkness. Whole-cell patch configuration was achieved and neuronal recordings were obtained at varying holding potentials as slices were illuminated with a blue LED pulse (a single pulse of 2 to 200 ms, or a train of 2 ms pulses at 10 to 100 Hz, power 11.5 mW) to elicit postsynaptic events. Where required, SR-95531 (2 µM; henceforth referred to as GABAzine), NBQX (10 µM) or CGP-52432 (10 µM) (all from Tocris, Bristol, UK) were washed into the perfusion bath via the perfusion pump.

### Immunohistochemistry and neuronal recovery

Following electrophysiological recordings, slices containing neurons which had been patched and filled with biocytin were processed as previously described (Wozny and Williams, 2011). Briefly, slices were fixed overnight in 4% paraformaldehyde (PFA) dissolved in 0.1 M sodium-based phosphate buffered saline (PBS). After fixation, slices were washed 3 x 5 minutes in 0.1 M PBS, and then incubated for 1 hour in a blocking solution consisting of 5% normal goat serum (NGS) and 1% Triton X-100. Slices were then allowed to incubate on a shaker at room temperature overnight in a primary antibody mixture containing 2.5% NGS and 1% Triton in PBS along with the required primary antibodies. Primary antibodies and dilutions used in this study were: mouse anti-PV (1/4000; Swant, Marly, Switzerland) and rabbit anti-GABA (1/200; Sigma-Aldrich, Dorset, UK). Upon completion of the primary incubation step, slices were washed 2 x 5 minutes in 0.1 M PBS and incubated for 2-3 hours in a secondary antibody cocktail containing the relevant secondary antibodies along with streptavidin (conjugated to Alex Fluor 488 or 647; 1/500 dilution; Life Technologies, Paisley, UK), in order to recover neurons which had been patched and filled with biocytin. The secondary antibodies used in this study were: donkey anti-mouse conjugated to Alexa Fluor 488 (1/500 dilution; Life Technologies, Paisley, UK), goat anti-rabbit conjugated to Alexa Fluor 555, 633 or 647 (1/500 dilution; Life Technologies, Paisley, UK) and supplemented with 1% Triton in PBS. Fluorophores excitable at differing wavelengths were implicated depending on whether the slice expressed YFP (Ai32 animals) or TdTomato (Ai9 animals) to minimize crosstalk. Where only neuronal recovery was required, slices were blocked as above and incubated in a solution containing streptavidin supplemented with 1% Triton in PBS. After secondary antibody incubation, slices were washed for 3 x 5 minutes in 0.1 M PBS and mounted on glass slides using Vectashield medium (containing DAPI as required, Vector Labs, Peterborough, UK) and cover-slipped.

### Double fluorescent in situ hybridization

Fresh, unfixed C57BL/6 mouse brains (N = 3) were embedded in tissue freezing medium (Leica Biosystems, Richmond, UK), frozen on a dry ice ethanol bath, sectioned at 20 μm on a cryostat, and then mounted onto Polysine Adhesion Slides (Thermo Fisher Scientific, Waltham, USA). RNA probe hybridization and subsequent washes were performed as described previously (Ansel et al., 2010). Fluorescein-labeled probes were detected using peroxidase-conjugated anti-fluorescein antibodies (Roche Diagnostics, Mannheim, Germany). Peroxidase activity was detected using Cy3-tyramide conjugate. After the detection, peroxidase activity was blocked by 100 mM glycine-HCl (pH 2.0) solution containing 0.1% Tween. Sections were washed with 0.1M Tris-HCl buffer (pH 7.5) containing 0.15M NaCl and 0.05% Tween and then blocked by 10% goat serum and 1% blocking reagent powder (Roche Diagnostics, Mannheim, Germany) in 0.1M Tris-HCl (pH 7.5) with 0.15M NaCl. Dioxygenin (DIG)-labeled probes were detected using peroxidase-conjugated anti-DIG antibodies (Roche Diagnostics, Mannheim, Germany). Peroxidase activity was detected using fluorescein-tyramide conjugate. Sections were counterstained with 4′,6-diamidino-2-phenylindole (DAPI) (Sigma-Aldrich, Munich, Germany).

### Tyramide conjugate synthesis

Fluorescein- and Cy3-tyramide conjugates were synthetized as described previously (Hopman et al., 1998). Briefly, the succinimidyl esters of fluorescein (Thermo Fisher Scientific, Schwerte, Germany) and Cy3 (GE Healthcare, Little Chalfont, United Kingdom) were coupled to tyramine (Sigma-Aldrich, Munich, Germany) in dimethylformamide (Carl Roth, Karlsruhe, Germany) adjusted to a pH of 7.0–8.0 with triethylamine (Sigma-Aldrich, Munich, Germany).

### Details of probes designed for in situ hybridization

DNA fragments used as template for RNA probe transcription were PCR-amplified from C57BL/6 mouse genomic DNA using the following primers:

VGAT: GACCTCGAGCTACCTGGGGTTGTTCCTCA, AATTAACCCTCACTAAAGGGACTAGTCGAAGTGTGGCACGTAGATG
VGLUT2: GACGAATTCTATTCGTTGGACCCATCACC, gacAATTAACCCTCACTAAAGGGCGGCCGCAGAAATTGCAATCCCCAAAC

PV:

Exon3: GACGAATTCCCTCTCCCCTGTCCTTCTTT, AATTAACCCTCACTAAAGGGCGGCCGCaTGGGAACTTTGGGTGCTATC
Exon4: GACGAATTCAGGTTCTGCCTGTGACCTTG, AATTAACCCTCACTAAAGGGCGGCCGCtAAGCTTTGACAGCCGCATAC
Exon5: GACGAATTCCTCCACTCTGGTGGCTGAA, AATTAACCCTCACTAAAGGGCGGCCGCTTTCTCTTTTCAGGTATTTTATCACA

Amplified DNA fragments were cloned into the pBSK backbone and transcribed with T3 RNA polymerase using DIG RNA Labeling Mix and Fluorescein RNA Labeling Mix according to the manufacturer protocol (all from Roche Diagnostics, Mannheim, Germany).

### Intracardial perfusion and serial sectioning

To prepare tissue for serial sectioning, Ndnf-IRES-Cre::Ai32 (N = 2; P25), Ndnf-IRES-Cre::Ai9 (N = 2; P21), PV-IRES-Cre::Ai32 (N = 2; P31-35), SOM-IRES-Cre::Ai9 (N = 2; P28), NPY-hrGFP (N = 2; P23) or C57BL/6 (N = 3; P21-22) mice were terminally anaesthetized by subcutaneous injection with an overdose cocktail of 50% lidocaine and 50% euthatal. Once anaesthetized sufficiently to be non-responsive to noxious tail and toe pinch stimuli, mice were perfused through the left ventricle with 0.1 M PBS followed by perfusion with 4% PFA dissolved in PBS. Brains were then removed and fixed overnight in 4% PFA in PBS, after which they were cryoprotected in a solution containing 30% (w/v) sucrose in PBS for storage until required for serial sectioning.

For these experiments, brains were embedded in OCT compound (VWR, Leicestershire, UK) and sectioned on a Leica SM2010 R microtome (Leica Biosystems, Newcastle-upon-Tyne, UK) at 60-80 µm. Upon completion of sectioning, slices were washed 3 x 5 min in 0.1 M PBS. Where further staining was required, this was carried out as above (see immunohistochemistry and neuronal recovery), however, 0.3% Triton X-100 was used in place of 1% and slices were incubated overnight in primary antibody cocktails to minimize tissue damage. Slices were then mounted using Vectashield medium (Vector Labs, Peterborough, UK) and cover-slipped.

### Image acquisition and neuronal reconstructions

For immunohistochemistry-stained sections and biocytin-filled neurons; mounted sections were scanned on either a Leica SP5 or SP8 confocal microscope, imaging z-stacks of each slice at 2-4 µm steps. Confocal laser excitation wavelengths (in nm) were 405, 488, 514, 552 and 503. Objectives used were 10X (dry), 20X (oil immersion), 40X (oil immersion) and 63X (oil immersion) for Leica SP5, or 10X (dry), 20X (dry) and 63X (oil immersion) for SP8. A zoom of up to 2X was applied as required to occasionally visualize soma in enhanced detail. Sections were scanned to ensure that all visible streptavidin-stained cells and their neurites were included in the z-stack. 3D reconstructions of neurons were carried out using NeuTube 3D reconstruction software(Feng et al., 2015).

For in situ hybridizations; the two-dimensional overview images were acquired with an Axio Examiner microscope with Axiocam 506 camera and LED Light Source Colibri 7 system using a Plan-Apochromat 20x/0.8 M27 objective (all from Carl Zeiss, Jena, Germany). The illumination wavelengths of 450-488 nm, 540-570 nm and 370-400 nm and emission filter wavelengths of 500-550 nm, 570-640 nm and 420-470 nm were used for Fluorescein, Cy3 and DAPI, respectively. The light source intensity was set to 40%, 50% and 20% for Fluorescein, Cy3 and DAPI, respectively. The three-dimensional images were acquired with a Zeiss LSM880 confocal laser scanning microscope using a Plan-Apochromat 20x/0.8 M27 objective (all from Carl Zeiss, Jena, Germany) with a 2x optical zoom. Laser wavelengths of 561 nm, 488 nm and 405 nm were used for Cy3, Fluorescein and DAPI, respectively. Stacks of 6 to 12 optical slices were captured with a z-step size of 0.785 μm.

### Data analysis

Analysis of electrophysiological recordings was carried out using Axograph X. Passive intrinsic properties were calculated as described above, while active intrinsic properties (action potential initial frequency, amplitude, rise-time and half-width) were calculated by subtracting the baseline and then using the event detection feature to analysis the first action potential elicited in response to a 50 pA depolarizing pulse. Neuronal spontaneous activity was classified as either bursting (a clearly distinctive behaviour), tonic or silent (where spontaneous action potential frequency was < 1 Hz; see Weiss and Veh, 2011). For optogenetically evoked events, peak size was measured at various holding potentials. For spontaneous current measurements, representative example traces of postsynaptic currents were first generated and currents were detected and measured using the event detection feature.

Image analysis was carried out using ImageJ. Confocal z-stacks were compressed onto a single image and brightness and contrast were occasionally adjusted to enhance cellular visualization. Cell counts were quantified using the cell-counter plugin. For these experiments, serial sections containing the whole habenula were imaged and analysed from one animal for each strain, and for remaining animals every second or third section was imaged and analysed to allow quantification of markers with fair representation of the habenular sub-nuclei. Images were then transferred to PowerPoint (Microsoft), where cells of interest were marked.

Graphs were generated and statistical analysis was performed using GraphPad Prism 5 (California, USA). Statistical tests used were: two-tailed unpaired t-test for single comparisons of passive physiological properties; one-way ANOVA with Tukey’s multiple comparison test for comparison of physiological properties between multiple groups; two-way ANOVA with Bonferroni’s multiple comparison for assessing relationship between input current and action potential discharge (fI-curves; fI-analysis), or Fishers’ exact test. Once graphs were generated, they were transferred to PowerPoint 2013 for formatting and assembly into figures. Statistical significance thresholds for all tests were: * *p* < 0.05; ** *p* < 0.01; and *** *p* < 0.001.

## Results

### Excitatory and inhibitory transmission within the lateral habenula

GABAergic signalling has previously been reported in the LHb (Lecca et al., 2016; Meye et al., 2016). To assess the balance of excitation and inhibition within the LHb, we simultaneously recorded spontaneous excitatory and inhibitory currents in LHb neurons using a low chloride intracellular recording solution (Fig. 1A; n = 10 neurons; N = 5 mice). For comparison, we recorded spontaneous events in L2/3 pyramidal neurons in the somatosensory cortex (n = 9; N = 4 mice). While the ratio of excitatory to inhibitory events was far larger in LHb neurons than in cortical neurons (Fig. 1B; 22.9 vs 2.3, respectively), we could clearly observe inhibitory currents with comparable frequency (Fig. 1C; 0.1 ± 0.0 Hz vs 0.5 ± 0.3 Hz, respectively; *p* = 0.23; two-tailed unpaired t-test), amplitude (Fig. 1C; 13.8 ± 1.3 pA vs 12.6 ± 1.1 pA, respectively; *p* = 0.49; two-tailed unpaired t-test) and kinetics (Fig. 1D; rise time 1.3 ± 0.2 ms vs 1.0 ± 0.1 ms, respectively; *p* = 0.11; half-width 2.1 ± 0.3 ms vs 1.7 ± 0.4 ms, respectively; *p* = 0.45; decay 16.1 ± 3.4 ms vs 13.4 ± 4.1 ms, respectively; *p* = 0.64; two-tailed unpaired t-test) to those in cortical neurons. Hence we asked whether these events could be mediated by similar inhibitory neurons as those in the neocortex.

**Figure 1:**
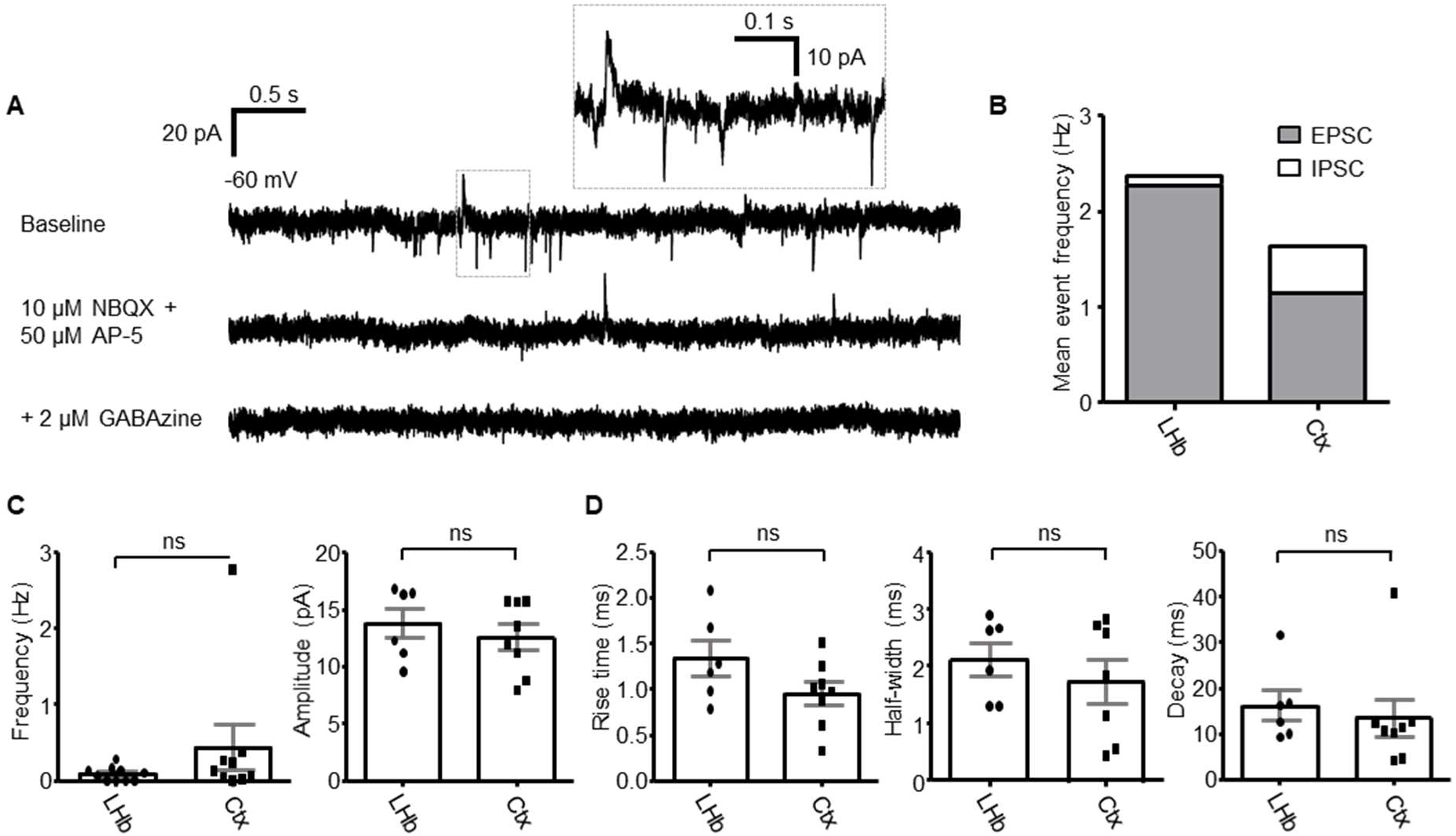
Spontaneous inhibitory currents within the lateral habenula are similar to those within the somatosensory cortex. **(A)** Example traces of spontaneous currents in an LHb neuron recorded in voltage clamp configuration. AMPA and NMDA-mediated excitatory currents and GABA_A_-mediated inward currents could be observed simultaneously using a low-chloride intracellular solution. **(B)** Relative frequency of excitatory vs inhibitory currents in both LHb neurons (n = 10 neurons from 5 mice) and somatosensory cortex L2/3 pyramidal neurons (n = 9 neurons from 4 mice). **(C)** Comparison of frequency (left) and amplitude (right) between inhibitory currents in LHb neurons and L2/3 pyramidal neurons. Data are mean ± SEM. **(D)** Comparison of kinetics (rise time, half-width and decay) between inhibitory currents in LHb neurons and L2/3 pyramidal neurons. Note for 4 LHb neurons, and 1 cortical neuron, no inhibitory currents were observed. For these neurons, frequency value was 0 Hz. However, as there were no measurable currents, these neurons have been excluded from analyses of current amplitude and kinetics.

### Characterisation of inhibitory sub-populations of LHb neurons

To address this question, we implemented the use of four well-characterized markers of inhibitory identity within the neocortex: PV (Tremblay et al., 2016), somatostatin (Tremblay et al., 2016), Ndnf (Abs et al., 2018; Tasic et al., 2018, 2016) and NPY (Overstreet-Wadiche and McBain, 2015). To test if these markers were also representative of inhibitory identity within the LHb, we used either transgenic mouse lines where fluorescent reporter proteins are expressed within neurons expressing the respective marker, or an antibody against the respective marker, and co-stained sections from these animals with an antibody against GABA.

We first made use of NPY-hrGFP mice, where GFP is expressed in NPY-positive neurons. Brains from these mice (N = 2) were sectioned in the coronal plane and imaged on a confocal microscope. In slices from NPY-hrGFP mice, while we did observe some sparse fibres in the LHb, we did not observe any NPY-positive somata (Fig. S1). Thus, we did not perform any further testing with NPY-hrGFP mice. We next crossed SOM-IRES-Cre and Ndnf-IRES-Cre mice to Ai9 (Madisen et al., 2010) reporter mice, so as to generate SOM-IRES-Cre::Ai9 (N = 2) and Ndnf-IRES-Cre::Ai9 offspring (N = 2), which expressed TdTomato in SOM-positive and Ndnf-positive neurons respectively (Fig. 2A). We could clearly observe TdTomato-expressing somata within the LHb in slices from both of these lines (Fig. 2B and D). Strikingly, the majority of SOM-positive neurons were clustered in the superior sub-nucleus of the LHb (Fig. 2Bb) (Andres et al., 1999), while Ndnf-positive neurons were mostly confined to the medial portion (Fig. 2D). However, in contrast to the neocortex (Fig. S2), both SOM-positive and Ndnf-positive LHb neurons were primarily non-GABAergic populations, as only a small sub-population of these (3.65 % and 14.3 % respectively) co-localised with GABA (Fig. 2C and E). Thus we conclude that both SOM-positive and Ndnf-positive LHb neurons are primarily non-GABAergic populations.

**Figure 2:**
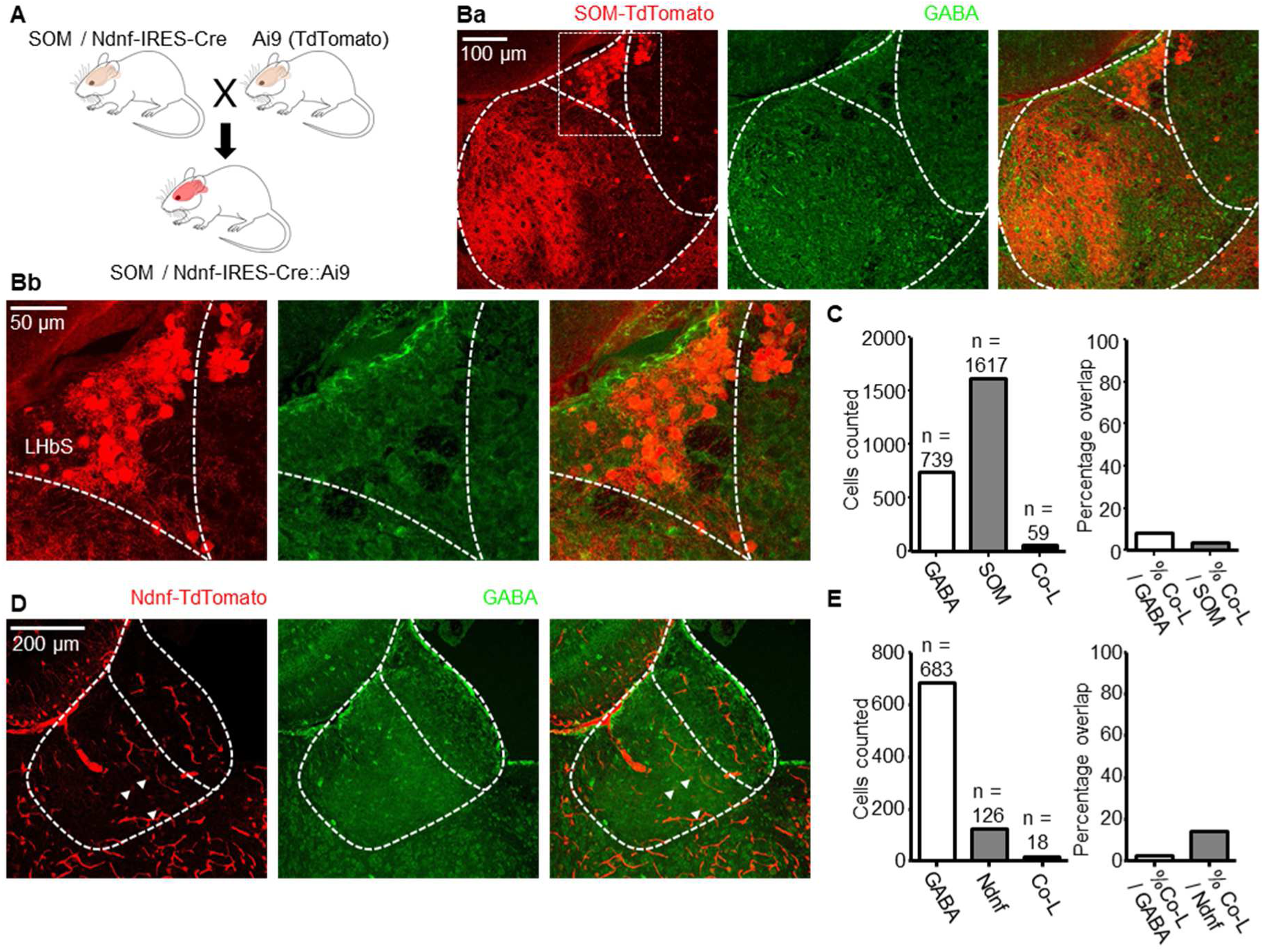
SOM and Ndnf-positive LHb neurons are primarily non-GABAergic populations. **(A)** Breeding scheme for generating SOM-IRES-Cre::Ai9 and Ndnf-IRES-Cre::Ai9 mice. **(Ba)** Confocal micrograph from SOM-IRES-Cre::Ai9 slices depicting SOM-positive neurons in the LHb. **(Bb)** Zoom of boxed region in (Ba) depicting SOM-positive LHb neurons localized within the superior sub-nucleus (LHbS) of the LHb. **(C)** Left: bar chart quantifying total number of GABA-immunoreactive, and SOM-positive neurons counted and the number which co-expressed both (N = 2 mice). Right: fraction of neurons expressing both markers as a percentage of GABA-immunoreactive neurons, and as a percentage of SOM-positive neurons. **(D)** Confocal micrograph as for (Ba), with Ndnf-postive neurons. Arrowheads indicate non-GABAergic Ndnf-positive neurons. **(E)** Quantification as for (C), with Ndnf-positive neurons.

We next stained slices from C57BL/6 mice (N = 3) with an antibody against PV. While PV-positive neurons were also clearly visible within the LHb (Fig. 3A and B), a similarly small fraction of these (8.8 %; N = 3 mice) were co-labelled with GABA (Fig. 3C). Interestingly, these formed two distinct clusters; one within the medial LHb (Fig. 3Aa) and one within the lateral LHb (Fig. 3Ab) along the rostral-caudal axis. Moreover, the GABAergic PV-positive neurons appeared to be exclusively confined to the lateral LHb (Fig. 3Ac and B), and were very brightly labelled with GABA (Fig. 3Ac), thus forming a sub-population of GABAergic neurons that was not observed in either the SOM or Ndnf lines. Despite this clear labelling however, background with the GABA antibody was relatively high (Figs. 2B and D, and 3A). We therefore also performed more sensitive in situ hybridizations with probes for PV and vesicular GABA transporter, VGAT (Fig. 3D; N = 3 mice); and PV and vesicular glutamate transporter 2, VGLUT2 (Fig. S3; N = 3 mice). Consistently, we could clearly observe PV and VGAT double-positive neurons within the lateral LHb (Fig. 3Db), while PV-positive neurons within the medial LHb were VGAT-negative (Fig. 3Dc), but VGLUT2-positive (Fig. S3B). Altogether, while these results indicate that the majority of PV-positive LHb neurons are non-GABAergic, they also suggest the existence of a unique sub-class of inhibitory PV-positive neurons located within the lateral LHb.

**Figure 3:**
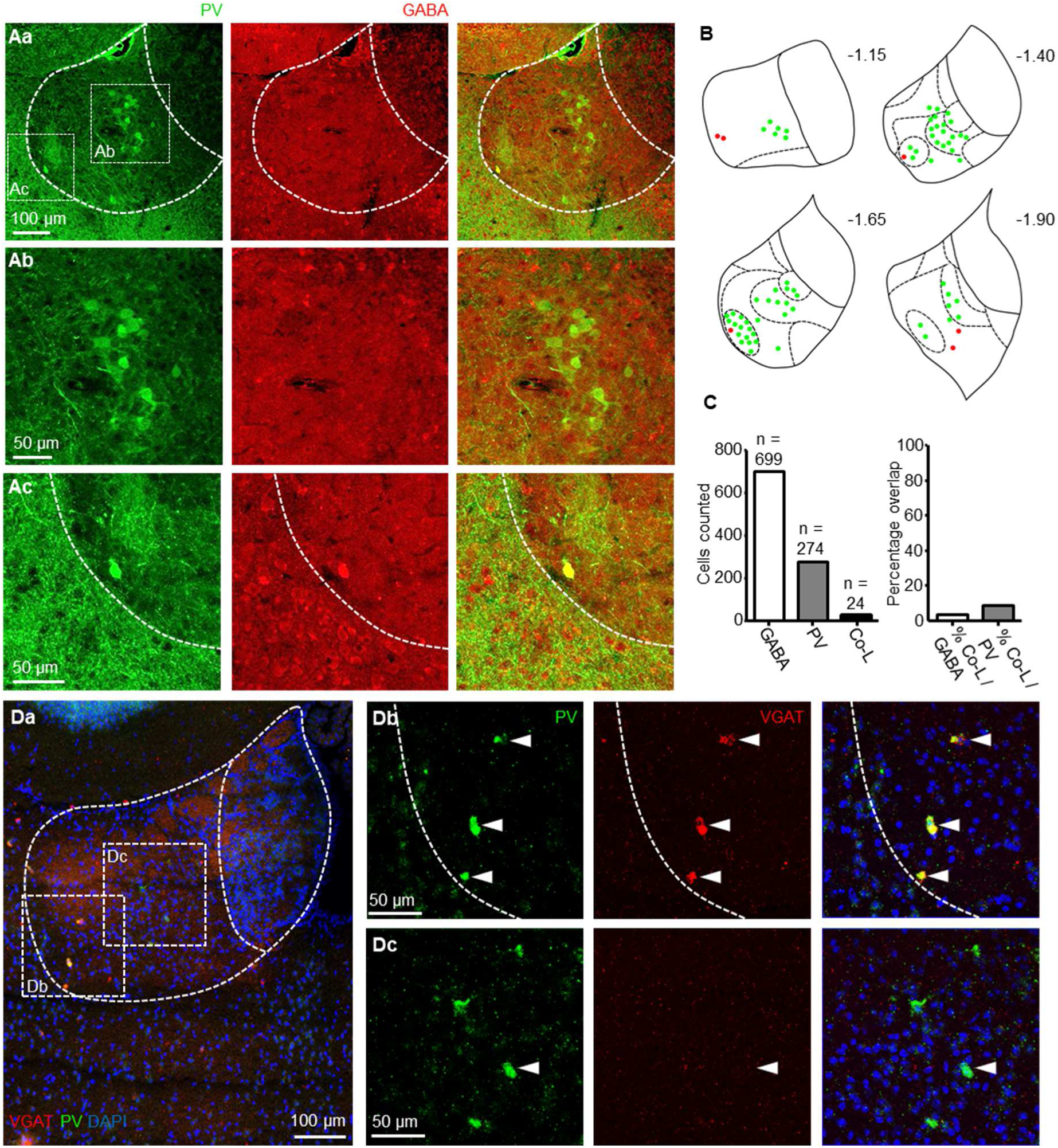
A sub-class of PV-positive neurons located within the lateral LHb are GABAergic. **(Aa)** 20x representative confocal micrographs displaying PV-immunoreactivity (left), GABA-immunoreactivity (middle) and merge of both (right) within the LHb. **(Ab)** Zoom of boxed region in (Aa) depicting PV-immunoreactive neurons which did not co-localise with GABA. **(Ac)** Zoom of boxed region in (Aa) depicting a PV-immunoreactive neuron which did co-localise with GABA. **(B)** Schematic illustrating location of PV-immunoreactive only, or PV / GABA co-labelled neurons throughout the LHb in the rostral-caudal plane from one mouse, in which every second 60 µm section was analysed. Sub-nuclear boundaries as defined by Andres et al., (1999) are indicated by dashed lines. Approximate rostral-caudal distances from Bregma (in mm) are indicated. **(C)** Bar graphs showing total number of GABA-immunoreactive, PV-immunoreactive and GABA / PV co-localising (Co-L) LHb neurons (left) and fractions of co-localising neurons as a percentage of total PV-immunoreactive and of GABA-immunoreactive neurons (right; N = 3 mice). **(Da)** 20X in situ hybridization overview image displaying the LHb co-stained with probes for PV and VGAT. **(Db)** Zoom of the left boxed region in (di) displaying VGAT-positive PV-positive neurons in the lateral LHb. **(Dc)** Zoom of the right boxed region in (Da) displaying VGAT-negative PV-positive neurons in the medial LHb.

### PV-positive and SOM-positive LHb neurons form physiologically distinct sub-classes

We next sought to further characterise Ndnf, PV and SOM-positive LHb neurons by assessing their physiological properties. We crossed each Cre-driver line with the Ai9 (Madisen et al., 2010) reporter line to generate Ndnf-IRES-Cre::Ai9 (N = 5), PV-IRES-Cre::Ai9 (N = 8) and SOM-IRES-Cre::Ai9 (N = 3) transgenic offspring (Fig. 4A), and used fluorescence-assisted patch-clamp recordings to record from TdTomato-expressing neurons in acute slices from each line (n = 29 Ndnf neurons; n = 19 PV neurons; n = 24 SOM neurons). We also recorded from a control sample of neurons from the general population of LHb neurons (n = 28 from 5 C57BL/6 mice), and compared passive physiological properties between all groups (Fig. 4B). Resting membrane potential was comparable between all groups (*p* = 0.23; one-way ANOVA); however clear differences could be observed between input resistances (*p* < 0.0001; one-way ANOVA). SOM-positive neurons had a far larger mean input resistance than any other group (1364.0 ± 111.3 vs 544.0 ± 53.3, 755.9 ± 76.5 and 366.2 ± 39.1 MΩ for general population, Ndnf and PV neurons respectively; *p* < 0.0001; Tukey’s multiple comparison test). Ndnf-positive neurons had the second largest input resistance, which was significantly greater than that of PV-positive neurons (*p* < 0.001; Tukey’s multiple comparison test), but not the general population (*p* > 0.05; Tukey’s multiple comparison test). Interestingly, both SOM-positive and Ndnf-positive neurons are generally located near the border with the medial habenula (Figs. 2, and 4D and N), where neurons are known to have very large input resistances (Choi et al., 2016), and as such these findings lend support to the recently proposed idea of an area of overlap between the lateral and medial habenulae (Wagner et al., 2016b).

**Figure 4:**
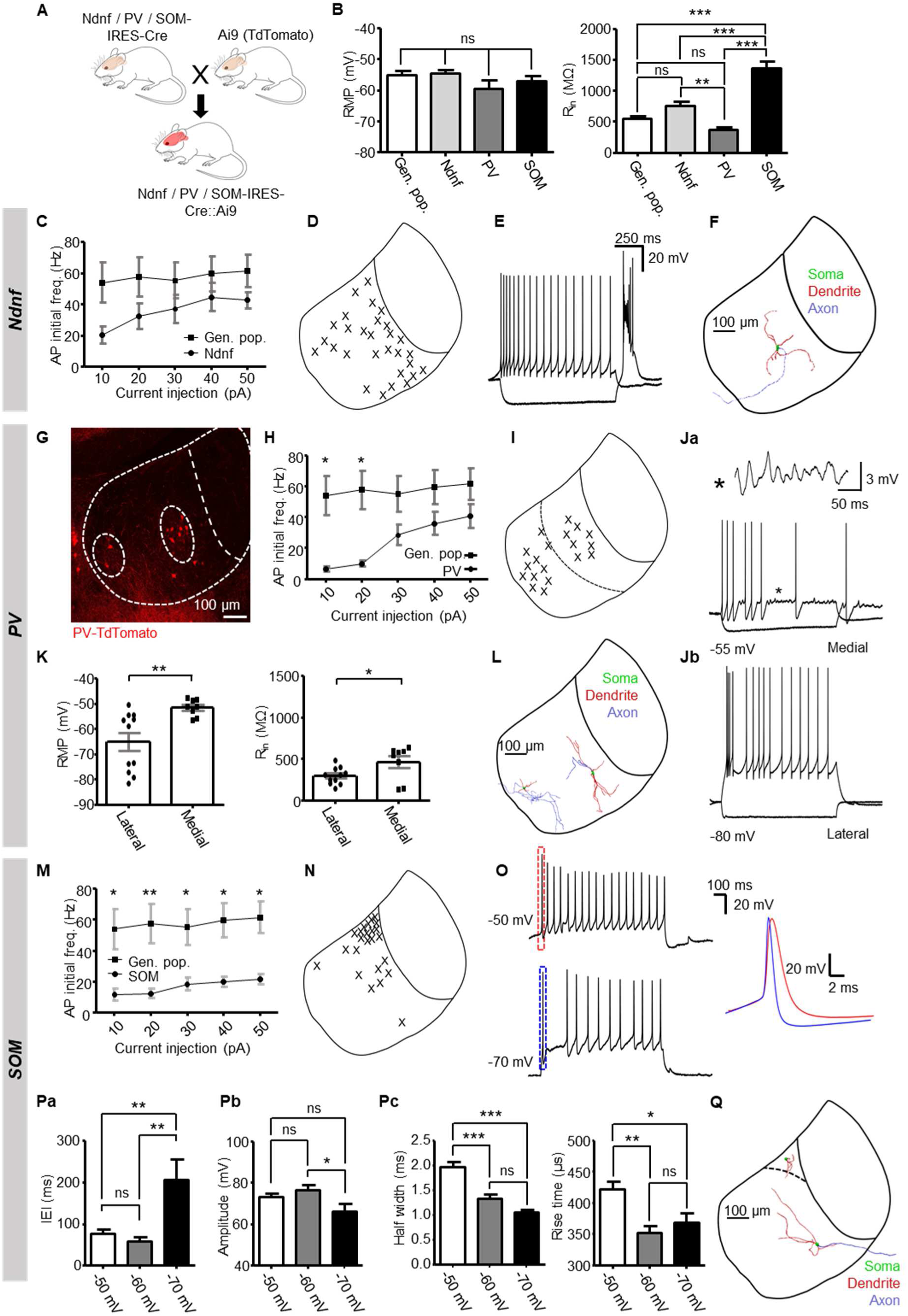
PV-positive and SOM-positive LHb neurons form physiologically distinct sub-classes. **(A)** Breeding scheme for generating Ndnf, PV and SOM-IRES-Cre::Ai9 transgenic mice. **(B)** Comparison of passive physiological properties between all groups. **(C)** Comparison of initial action potential firing frequency versus current injection for both Ndnf-positive LHb neurons (n = 23; N = 5 mice) and the general population of LHb neurons (n = 28; N = 5 mice) recorded in C57 mice. Data are mean ± SEM. **(D)** Schematic illustrating location of patched neurons throughout the habenular complex. In this case, all neurons recorded have been compressed onto one schematic to allow clear visualization of neurons recorded near the border with the medial habenula vs throughout the rest of the LHb. **(E)** Example traces from an Ndnf-positive neuron in response to current injection. Current steps: −50 pA and 50 pA; 1 s duration. (**F**) Example morphological reconstruction from an Ndnf-positive neuron. All reconstructed neurons (n = 5) displayed 4-6 primary dendrites and a single unbranching axon. **(G)** Confocal micrograph depicting localization of PV-positive neurons within the LHb. The central and oval sub-nucleus approximate boundaries are indicated by dashed white lines. **(H)** As for (C), PV-positive neurons (n = 18; N = 8 mice). **(I)** As for (D), for PV-positive neurons. In this case, all neurons recorded have been compressed onto one schematic to allow clear visualization of neurons recorded in the lateral vs medial LHb. **(J)** Example traces from PV-positive neurons in response to current injection. (Ja): traces from a neuron in the medial LHb. Asterisk indicates a section from the recording displaying characteristic sub-threshold voltage oscillations. (Jb): traces from a neuron in the lateral LHb. Current steps: −50 pA and 50 pA; 1 s duration. **(K)** Comparison of passive properties between PV positive neurons in the lateral LHb (n = 11) and medial LHb (n = 8). Data are mean ± SEM. **(L)** Example reconstructions from two PV-positive neurons. Left: a neuron with an extensively-branching axon (observed in 1 of 10 reconstructions). Right: a representative reconstruction of neuron which displays an unbranching axon (observed in 9 of 10 reconstructions). **(M)** As for (C), for SOM-positive LHb neurons (n = 22; N = 3 mice). **(N)** As for (D), in for SOM-positive neurons. **(O)** Example traces from one neuron when held at −50 mV and −70 mV in response to a 1 s injection of 50 pA current. Zoom: comparison of the first induced action potential at each holding potential. **(Pa)** Inter-event interval (IEI) between the first and second action potential induced in response to depolarising current injection (20-150 pA; 1 s) for neurons tested at different holding potentials (n = 11). **(Pb)** Comparison of amplitude of the first (‘early spike’) action potential induced in response to depolarising current injection (20-150 pA; 1 s) for neurons tested at different holding potentials (n = 11). **(Pc)** Comparison of kinetics (rise time and half-width) of the first action potential induced in response to depolarising current injection (20-150 pA; 1 s) for neurons tested at different holding potentials (n = 11). **(Q)** Example reconstructions from a SOM-positive neuron within the superior sub-nucleus (top; n = 6), and from another SOM-positive neuron out with the superior sub-nucleus (bottom; n = 4). Note that the neuron in the superior sub-nucleus displays short stubby dendrites, while the other neuron displays a more conventional LHb neuronal morphology.

We next assessed the active physiological properties of each class of neuron by injection of a series of current steps. In two Ndnf-positive cells, action potential discharge could not be elicited upon current injection and hence we assumed these to be glial cells and excluded from further analysis. In the remainder we did observe an overall difference in the relationship between input current and action potential discharge frequency (Fig. 4C; *p* = 0.0008; two-way ANOVA; n = 23 neurons tested) in comparison to the general population (n = 28 neurons tested). Otherwise, physiological properties of Ndnf-positive LHb neurons were largely consistent with previously described LHb neuronal physiologies (Kim and Chang, 2005; Weiss and Veh, 2011; Yang et al., 2018) in that almost all (n = 22 from 24 neurons tested) displayed rebound action potential discharge upon hyperpolarizing current injection, and a combination of tonic and bursting action potential discharge upon depolarizing current injection (Fig. 4E). We also reconstructed a small subset of these neurons (n = 5) and observed that all neurons reconstructed exhibited 4-6 primary dendrites, and a long unbranching axon (Fig. 4F), again largely consistent with previous reports of generic lateral habenular neurons (Kim and Chang, 2005; Weiss and Veh, 2011), and as such we concluded that Ndnf expression was not confined to any particular sub-population of neuron within the LHb.

Consistent with our histological data, most PV-positive neurons were clustered in either the medial or lateral LHb (Fig. 4G and I). There was also a clear reduction in the firing frequency of the first induced action potential in response to depolarising current injection in comparison to neurons in the general population (Fig. 4H; *p* < 0.0001; two-way ANOVA). This was as a result of the fact that only a minority (4 of 19) of PV-positive neurons exhibited any kind of high-frequency bursting behaviour (Fig. 4Ja), and this was only observed upon larger current injections (Fig. 4H). This was a striking observation as high-frequency bursting has long been considered a hallmark physiological phenotype of most LHb neurons (Weiss and Veh, 2011; Wilcox et al., 1988; Yang et al., 2018). Furthermore, the clusters of PV-positive neurons in the medial and lateral LHb could clearly be differentiated based on their physiological profile (Fig. 4I, J and K). PV-positive neurons in the medial LHb frequently exhibited sub-threshold voltage oscillations (6 of 8 medial LHb neurons) while their counterparts in the lateral LHb did so very rarely (1 of 11 lateral LHb neurons, *p* = 0.006; Fishers exact test; Fig. 4J). Moreover, a second distinctive population of PV-positive neurons appeared in the lateral LHb, identifiable by their hyperpolarized resting membrane potential (5 of 11 lateral LHb neurons; −77.0 ± 1.3 vs −59.4 ± 2.6 mV for all 19 PV-positive neurons; *p* = 0.001; two-tailed unpaired t-test) and lack of rebound action potential discharge (Fig. 4Jb and K), a hallmark phenotype of LHb neurons (Chang and Kim, 2004; Kim and Chang, 2005; Weiss and Veh, 2011). We also performed morphological reconstruction of these neurons (Fig. 4L). Ten neurons were sufficiently reconstructed to visualize a prolonged section of the axon. In nine of these, this was an unbranching axon, possibly indicative of a projection neuron (Fig. 4L). In the one remaining neuron, we did observe the axon to branch locally and extensively, suggesting this to be a locally-targeting neuron. Taken together, these results indicate that PV-positive LHb neurons form multiple distinct sub-populations based on physiological profile and location within the LHb.

When recording active properties of SOM-positive neurons, a striking pattern quickly emerged. These neurons had a far lower firing frequency than the general population, regardless of input current (Fig. 4M; two-way ANOVA; p < 0.0001; and Bonferroni’s multiple comparison test; *p* < 0.05), and many of these neurons (15 of 24) displayed a prominent ‘early spike’ upon depolarising current injection (Fig. 4O; defined as spiking latency < 20 ms after current injection; current injection 20-150 pA, 1 s). Furthermore, we observed that this spike became more pronounced upon hyperpolarisation (Fig. 4O and P; tested in 11 neurons). Upon hyperpolarisation as far as −70 mV, we could observe a greater interval between this spike and the next spike in the train (Fig. 4Pa; *p* = 0.0014; one-way ANOVA), and that this early spike had faster kinetics than when depolarized (Fig. 4O and Pc; rise time *p* = 0.0011; half-width *p* < 0.0001; two-way ANOVA). Additionally, we reconstructed a subset of these neurons (Fig. 4Q; n = 10), and observed that of those in the superior sub-nucleus (n = 6), 5 of these displayed the short stubby dendrites associated with medial habenular neurons (Kim and Chang, 2005), while those outside of the superior sub-nucleus (n = 4) had the elongated dendrites known to be far more conventional of LHb neurons (Kim and Chang, 2005; Weiss and Veh, 2011). Membrane potential-mediated change in firing modality is a hallmark physiological characteristic of LHb neurons (Weiss and Veh, 2011; Wilcox et al., 1988; Yang et al., 2018). Yet these SOM-positive neurons discharge only one action potential as opposed to the bursting discharge commonly displayed by LHb neurons (Weiss and Veh, 2011; Wilcox et al., 1988; Yang et al., 2018), and then continue to discharge action potentials in a tonic train akin to that described in the medial habenula (Kim and Chang, 2005). These neurons also have huge input resistances (Fig. 4B) and similar morphological properties (Fig. 4Q) more comparable to that of medial habenular neurons. Thus, while we conclude that SOM-positive neurons are not GABAergic (Fig. 2), we provide physiological evidence for a sub-class of habenular neurons in the superior sub-nucleus (Figs. 2A and 4N) which possess intermediate characteristics of neurons from both the lateral and medial habenula, adding support to transcriptomic data proposing the existence of an area of overlap between the two regions (Wagner et al., 2016b).

### PV-positive and SOM-positive neurons provide inhibitory input to the LHb

Our data indicates that while some PV-positive LHb neurons are inhibitory (Fig. 3), Ndnf, PV and SOM-positive LHb neurons are likely not primarily inhibitory populations as they are known to be in the neocortex (Tasic et al., 2016; Tremblay et al., 2016). Additionally, previous studies have reported excitatory PV-positive neurons which project to the LHb (Knowland et al., 2017; Wallace et al., 2017) and GABA / glutamate co-releasing SOM-positive neurons which also target the LHb (Lazaridis et al., 2019; Wallace et al., 2017). Considering this information, we next asked whether each of these classes of neurons mediate primarily excitatory or inhibitory transmission within the LHb.

We therefore crossed mice from each Cre-driver line to Ai32 (Madisen et al., 2012) reporter mice, to generate offspring that express ChR2 and eYFP in Ndnf, PV or SOM-positive neurons respectively (Fig. 5Aa). We cut acute slices from these mice and recorded postsynaptic potentials in LHb neurons (Fig. 5Ab) in response to photostimulation. In slices from Ndnf-IRES-Cre::Ai32 mice (N = 6), most responsive neurons (n = 5 of 6, from 21 tested; Fig. 5B) displayed a solely excitatory postsynaptic potential (EPSP), while one neuron displayed a postsynaptic potential with both an NBQX-sensitive excitatory component and a GABAzine-sensitive inhibitory component (Fig. 5Ca and Cb). These responses were spread fairly evenly throughout the LHb (Fig. 5D), consistent with confocal imaging of serial coronal sections from Ndnf-IRES-Cre::Ai32 mice (N = 2), in that both Ndnf-positive fibres and somata were dispersed evenly throughout the LHb (Fig. S4). Taken together with our histological and physiological data (Fig. 2 and 4), we conclude that Ndnf is not expressed selectively by inhibitory neurons within, or projecting to the LHb, and thus carried out no further testing on Ndnf neurons; in clear contrast to Ndnf-positive neocortical neurons which mediate slow inhibitory signalling (Fig. S5).

**Figure 5:**
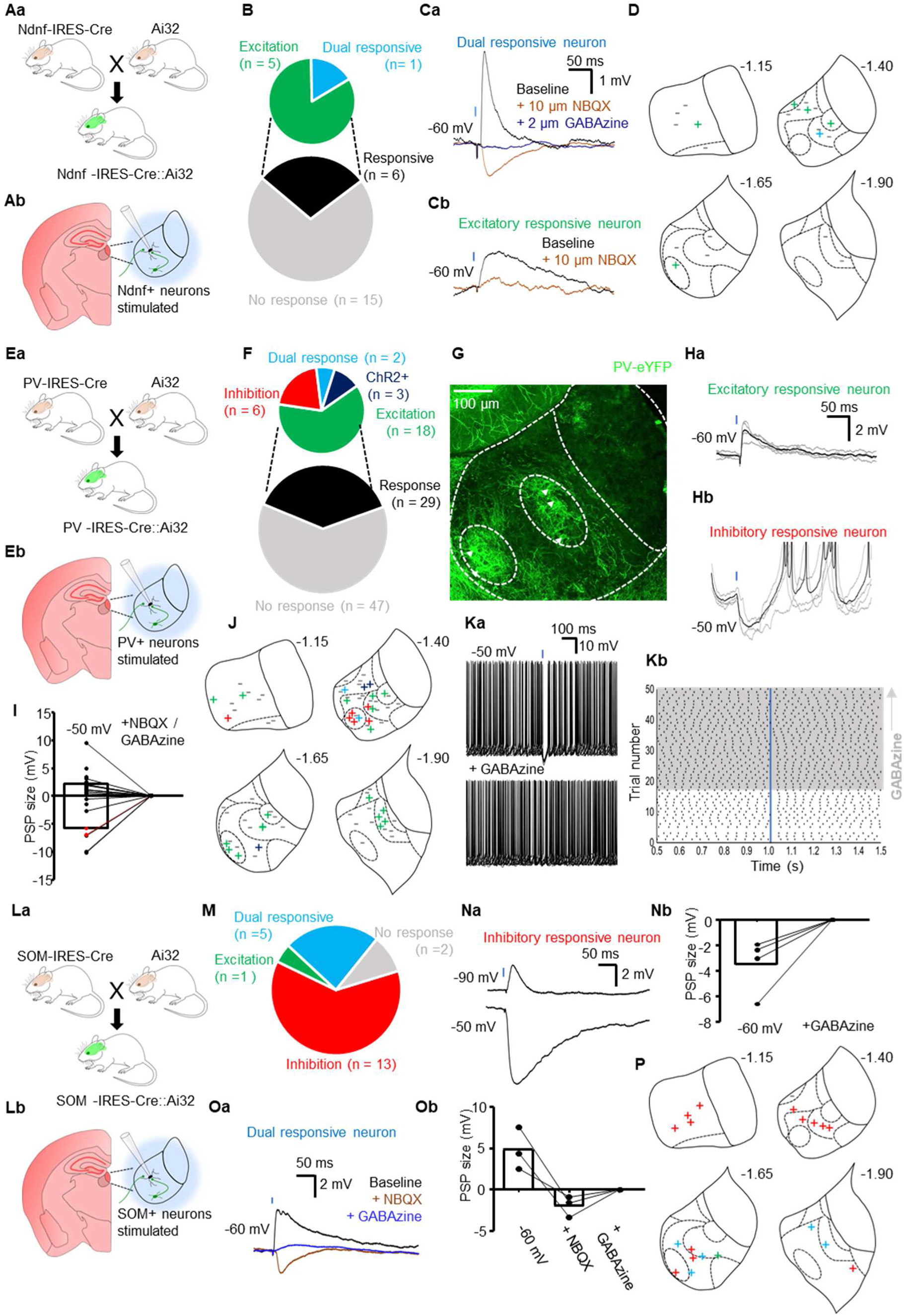
Optogenetic dissection of signalling pathways mediated by Ndnf, PV and SOM-positive neurons within the LHb. **(Aa)** Schematic illustrating breeding scheme for generating Ndnf, PV or SOM-IRES-Cre::Ai32 mice. **(Ab)** Schematic of recording scheme. LHb neurons were patched and blue light was used to stimulate both local and long-distance Ndnf, PV or SOM-positive neurons. **(B)** Pie chart quantifying fraction of neurons responsive to photostimulation, and the nature of those responses, in slices from Ndnf-IRES-Cre::Ai32 mice. **(Ca)** Example traces from a neuron in which photostimulation elicited a response with both an NBQX-sensitive excitatory component, and a GABAzine-sensitive inhibitory component and **(Cb)** from a neuron in which photostimulation elicited an excitatory response. **(D)** Schematic illustrating location of patched neurons within the habenular complex, projected rostrally (top left) through caudally (bottom right). Non-responsive neurons are indicated by **–** while responsive neurons are indicated by: **+** EPSP only; **+** dual response. Sub-nuclear boundaries as defined by Andres et al., (1999) are indicated by dashed lines. Approximate rostral-caudal distances from Bregma (in mm) are indicated. **(E)** As for (B), in PV-IRES-Cre::Ai32 slices. **(F)** Representative confocal micrograph depicting location of both PV-positive somata and fibres within the LHb. Arrowheads indicate PV-positive somata. **(Ga)** Example traces from one neuron in which light stimulation elicited an EPSP, and **(Gb)** one in which it elicited an IPSP (bottom). Traces are averages of multiple traces superimposed over the traces from which the average was derived. Blue square denotes 2 ms light pulse. **(H)** Before-after plot showing PSP size for neurons responsive to light which exhibited EPSPs (n = 16; note 2 neurons did not display a measurable EPSP at −50 mV) and IPSPs (n = 6) at - 50 mV and with application of either GABAzine (2 µM) or NBQX (10 µM) to abolish the PSP. Two neurons which displayed both excitatory and inhibitory components are highlighted in red. One of these neurons was tested pharmacologically, where both GABAzine and NBQX were applied to fully abolish the PSP (red line). **(I)** As for (D), in PV-IRES-Cre::Ai32 slices. Non-responsive neurons are indicated by **–** while responsive neurons are indicated by: **+** EPSP only; **+** IPSP only; **+** dual response consisting of both an excitatory and inhibitory component and; **+** ChR2-expressing neuron in which an action potential not sensitive to NBQX was triggered upon photostimulation. **(Ja)** Example traces overlaid from a neuron in which light-stimulation was sufficient to inhibit spontaneous action potential firing (top), and which was completely blocked in the presence of GABAzine (bottom). **(Jb)** Raster plot of (Ja). Grey area denotes presence of GABAzine. Blue bar denotes light 2 ms light stimulation. **(K)** As for (B), in SOM-IRES-Cre::Ai32 slices. **(La)** Example traces from a neuron at two different holding potentials in which photostimulation elicited an inhibitory postsynaptic potential. **(Lb)** Before-after plot from neurons receiving inhibitory input in which the IPSP was pharmacologically blocked (n = 4 neurons). **(Ma)** Example traces from a neuron in which photostimulation elicited co-release of both GABA and glutamate. **(Mb)** Before-after plot from neurons receiving GABA / glutamate co-releasing input in which both excitatory and inhibitory components of the postsynaptic potential were pharmacologically blocked (n = 3 neurons). **(N)** As for (D), in SOM-IRES-Cre::Ai32 slices.

In slices from PV-IRES-Cre::Ai32 transgenic mice imaged in the rostral-caudal axis (N = 2), we could visualize both PV-positive somata and fibres within the LHb (Figs. 5F and S6). As with the staining data (Fig. 3), and data from the PV-IRES-Cre::Ai9 line (Fig. 4G), PV-positive somata were mostly confined to two clusters in either the medial or lateral LHb (Figs. 5F and S6). These clusters also appeared densely enriched with fibres, and appeared to roughly correlate to the previously described central sub-nucleus of the medial LHb, or oval sub-nucleus of the lateral LHb (Andres et al., 1999). While this dense enrichment of fibres could be processes from local PV-positive neurons, we speculated that they may also be from upstream PV-positive projection neurons, as these are known to specifically target the oval sub-nucleus of the LHb (Wallace et al., 2017).

In acute slices from these mice (N = 19), photostimulation-induced postsynaptic potentials were observed in 29 of 76 (38.2 %) neurons recorded (Fig. 5E and G-I). The majority of these (18 of 29; 62.1 %) displayed a solely excitatory response (Fig. 5Ga). However, solely inhibitory responses were also observed with relative frequency (6 of 29 responsive cells; 20.7%; Fig. 5Gb) while on two occasions, we recorded postsynaptic potentials consisting of both an excitatory and inhibitory component (Fig. 5E). Furthermore, inhibitory postsynaptic potentials (IPSPs) were comparably larger than excitatory postsynaptic potentials (EPSPs) (mean 5.6 ± 1.3 vs 2.3 ± 0.6 mV, respectively; *p* = 0.013; unpaired t-test; Fig. 5H). In three of these neurons, the induced IPSP was sufficiently large to momentarily silence spontaneous action potential discharge of the recorded neuron (Fig. 5J) in a manner which could be blocked by application of GABAzine. In contrast, we report that input PV-positive neurons mediate both excitatory and inhibitory input to the LHb.

In SOM-IRES-Cre::Ai32 slices (N = 5 mice), we observed solely inhibitory responses in over half of the neurons tested (Fig. 5K and La; n = 13 of 21 neurons), confirmed by complete blockade upon GABAzine application (Fig. 5Lb; n = 4 neurons) We also frequently observed postsynaptic potentials consisting of both an excitatory and inhibitory component (Fig. 5K and M; n = 5 of 21 neurons); most likely arising from those previously described GABA / glutamate co-releasing neurons in the entopeduncular nucleus (Lazaridis et al., 2019; Wallace et al., 2017), which appeared to be specifically confined to the caudal portion of the LHb (Fig. 5N). Therefore we conclude that input to the LHb from SOM-positive neurons is primarily inhibitory.

### PV-positive LHb neurons mediate local inhibitory transmission

The above data (Fig. 5) shows that PV-positive neurons provide both excitatory and inhibitory input to the LHb, while SOM-positive neurons provide inhibitory input, and also co-release GABA and glutamate. Previous work can explain the origin of these excitatory (Knowland et al., 2017; Wallace et al., 2017), and co-releasing (Lazaridis et al., 2019; Wallace et al., 2017) inputs. So where then do these inhibitory inputs arise from?

Our histological data (Figs. 2 and 2) suggests that PV-positive inhibitory neurons, but not SOM-positive neurons, exist within the LHb. To verify this, and that these neurons make local synaptic connections, we used a viral approach. We injected PV-IRES-Cre mice and SOM-IRES-Cre mice with Cre-dependent AAVs encoding ChR2 and eYFP directly into the LHb to drive expression of ChR2 and eYFP in local PV and SOM-positive neurons (Figs. 6Aa and S7Aa). For PV-IRES-Cre mice, we injected an AAV containing either capsid protein 9 (N = 6), or a hybrid of capsid proteins 1 and 2 (N = 2). For SOM-IRES-Cre mice (N = 4) we used the serotype 9 virion. We then created acute slices from these mice and recorded postsynaptic events while photostimulating (Figs. 6Ab and S7Ab).

At least two weeks post-injection, we could observe robust eYFP expression in the LHb in both PV-IRES-Cre (Fig. 6B) and SOM-IRES-Cre (Fig. S7B) mice. Upon photostimulation of PV neurons, we could occasionally observe GABAzine-sensitive IPSP’s (Fig. 6C and D; n = 5 of 52 neurons tested) which were confined to the lateral portion of the LHb (Fig. 6E). Interestingly, we also observed one NBQX-sensitive excitatory response, and one response featuring both a GABAergic and glutamatergic component (Fig. 6C), possibly indicative that glutamatergic PV-positive neurons (Fig. S3) make local contacts. In the case of SOM-positive neurons, as expected we observed no synaptic responses (Fig. S7C and D), hence confirming that SOM-positive neurons do not make local inhibitory or excitatory contacts. Altogether, these data hence show that some PV-positive, but not SOM-positive, LHb neurons are locally-targeting inhibitory neurons.

**Figure 6:**
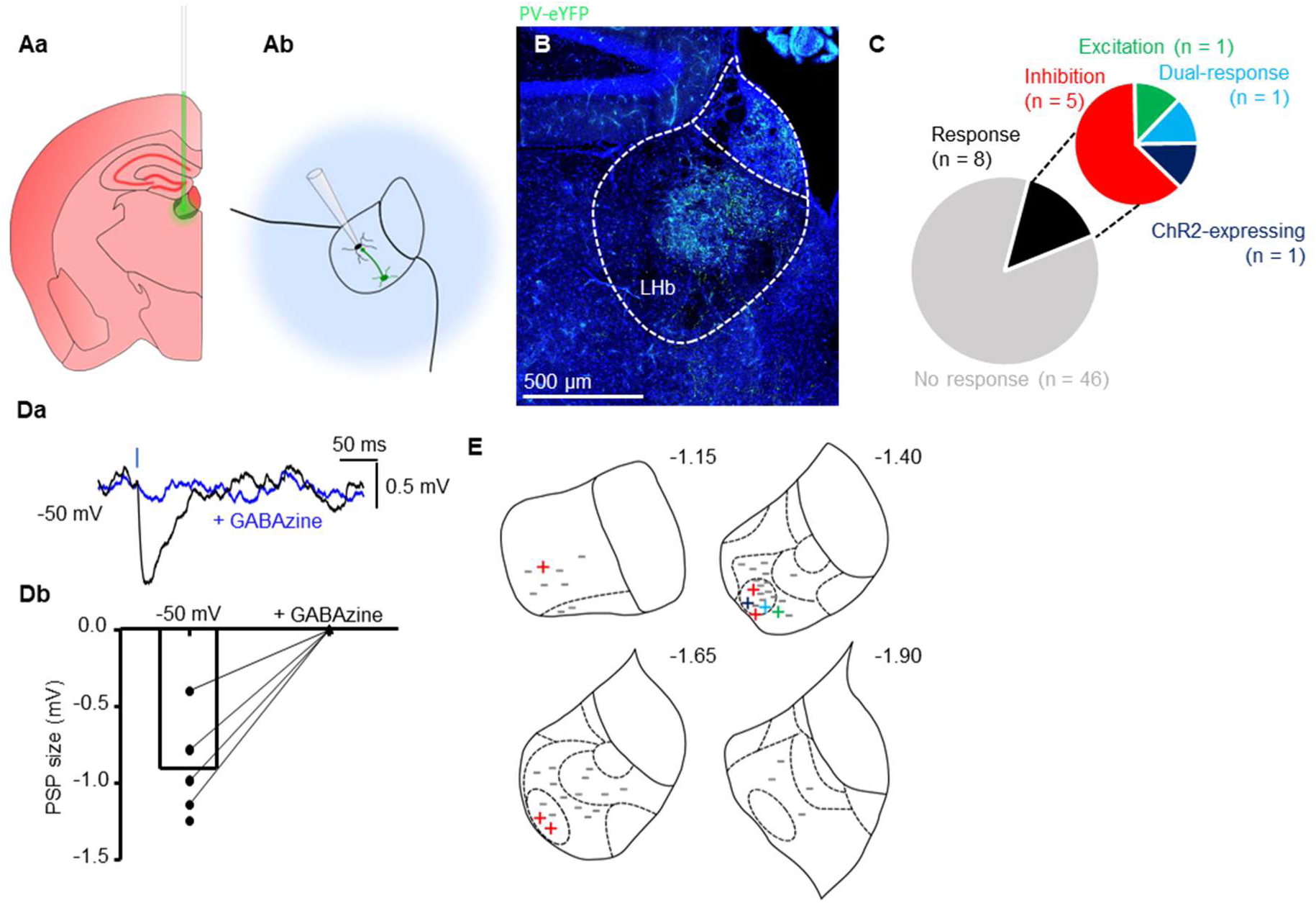
PV-positive neurons mediate local inhibitory signalling within the LHb. **(Aa)** Schematic illustrating stereotaxic injection protocol of AAV9 or AAV1/2 into the LHb of PV-IRES-Cre mice (N = 8). **(Ab)** Schematic illustrating electrophysiology recording protocol for LHb neurons following stereotaxic viral injection. Transduced neurons are photostimulated while recording from nearby LHb neurons. **(B)** Confocal micrograph depicting PV-positive LHb neurons transduced to express eYFP following viral injection. **(C)** Pie chart quantifying fraction of neurons responsive to photostimulation of PV-positive neurons. **(Da)** Example traces from one neuron in which a GABAzine-sensitive IPSP could be elicited following photostimulation. Blue bar denotes 2 ms photostimulation. **(Db)** Before-after plot of the IPSP amplitude from 5 different neurons at −50 mV before and after application of GABAzine (4 neurons). **(E)** Schematic illustrating location of patched neurons within the habenular complex, projected rostrally (top left) through caudally (bottom right). Non-responsive neurons are indicated by **–** while responsive neurons are indicated by: **+** EPSP only; **+** IPSP only; **+** dual response consisting of both an excitatory and inhibitory component and; **+** ChR2-expressing neuron in which an action potential not sensitive to NBQX was triggered upon photostimulation. Sub-nuclear boundaries as defined by Andres et al., (1999) are indicated by dashed lines. Approximate rostral-caudal distances from Bregma (in mm) are indicated.

### Distinct extrinsic inhibitory inputs to the LHb arise from PV-positive neurons in the medial dorsal thalamus and SOM-positive neurons in the ventral pallidum

If not local, then the large inhibitory responses we observed mediated by SOM-positive neurons (Fig. 5K and L) remained unaccounted for. We hence referred to the Allen Brain Atlas, and by searching for both SOM and GAD65/67 expression in the main afferent input regions to the LHb (Herkenham and Nauta, 1977), we observed that the ventral pallidum (VP) stood out as a region enriched with both SOM-positive and GAD-positive neurons, and speculated that this area may be the origin of the observed input.

Interestingly, recent work has shown that the lateral geniculate nucleus of the thalamus projects to and inhibits the LHb (Huang et al., 2019), and as such we speculated on the possibility that other thalamic neurons could also provide inhibitory input. Indeed, double in-situ hybridizations for PV and VGAT showed neurons positive for both these markers in the MDT (Fig. S8), and we therefore sought to address if both the MDT and VP were providing inhibitory input to the LHb.

We performed stereotaxic injection of Cre-dependent AAV9 encoding ChR2 and eYFP into the MDT of PV-IRES-Cre mice (Fig. 7A; N = 6), and into the VP of SOM-IRES-Cre mice (Fig. 7G; N = 3). By targeting injections to the ventral MDT, we could confine injections to this region without infecting the LHb (Fig. 7Aa and D). We recorded from LHb neurons (n = 47) while photostimulating MDT PV-positive neurons and observed inhibitory events in seven neurons (Fig. 7B and C; note in three these were only visible when the neuron was strongly depolarized). Consistently, upon post-hoc confocal imaging, we could visualize fibres which appeared to be projecting dorsally from neuronal somata located in the MDT to the LHb (Fig. 7D). Strikingly, and consistent with our observation of fibre enrichment (Fig. S5), these fibres appeared to be exclusively targeting the lateral LHb; particularly the oval sub-nucleus (Andres et al., 1999) where all responsive neurons were recorded (Fig. 7E). We also filled PV-positive MDT neurons (n = 8) with biocytin in slices from PV-IRES-Cre::Ai9 mice (N = 2), and upon reconstruction could observe fibres penetrating the LHb in 5 of 8 neurons (Fig. 7F).

**Figure 7:**
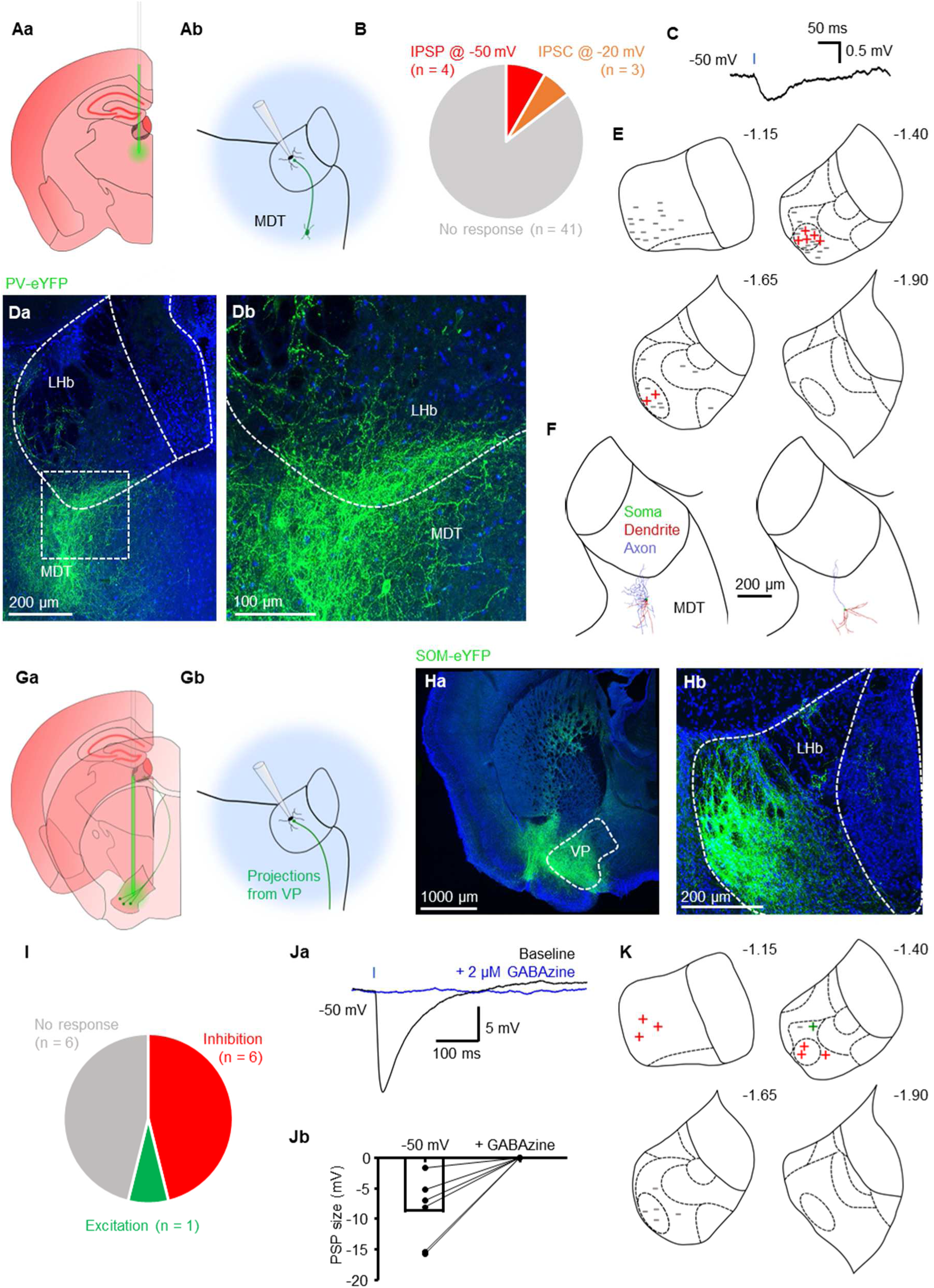
Distinct extrinsic inhibitory inputs to the LHb from the MDT and VP. **(Aa)** Schematic illustrating stereotaxic injection protocol of AAV9 into the MDT of PV-IRES-Cre mice (N = 6). **(Ab)** Schematic illustrating electrophysiology recording protocol for LHb neurons following stereotaxic viral injection. Transduced PV-positive neurons are photostimulated while recording from nearby LHb neurons. **(B)** Pie chart quantifying fraction of neurons responsive to photostimulation. **(C)** Example traces from one neuron in which an IPSP could be elicited following photostimulation. Blue bar denotes 2 ms photostimulation. **(Da)** Confocal micrograph depicting eYFP expression within the MDT following stereotaxic viral injection. **(Db)** Zoom of the outlined region in (Da). Note the presence of fibres originating from neurons in the MDT which penetrate the LHb. **(E)** Schematic illustrating location of patched neurons throughout the habenular complex. All neurons responsive to photostimulation were located in the oval sub-nucleus of the lateral habenula. Sub-nuclear boundaries as defined by Andres et al., (1999) are indicated by dashed lines. Approximate rostral-caudal distances from Bregma (in mm) are indicated. **(F)** Example reconstructions from 2 PV-positive MDT neurons which had axonal fibres projecting to the LHb. **(Ga)** Schematic illustrating stereotaxic injection protocol of AAV9 into the VP of SOM-IRES-Cre mice (N = 3). **(Gb)** Schematic illustrating electrophysiology recording protocol for LHb neurons following stereotaxic viral injection. Transduced SOM-positive neurons are photostimulated while recording from postsynaptic LHb neurons. **(Ha)** Confocal micrograph depicting transduced neurons following injection into the VP. **(Hb)** Confocal micrograph depicting terminals from transduced VP neurons in the LHb. **(I)** As for (B), in SOM-IRES-Cre slices. **(Ja)** Example traces from one neuron in which a GABAzine-sensitive IPSP could be elicited following photostimulation. Blue bar denotes 2 ms photostimulation. Traces are averages taken from multiple sweeps. **(Jb)** Before-after plot showing amplitude of photostimulation-induced IPSPs before and after application of GABAzine (n = 6 neurons). **(K)** As for (E), in SOM-IRES-Cre slices.

More than three weeks post-injection in SOM-IRES-Cre mice (Fig. 7G; N = 3), we could clearly see virally-transduced eYFP-expressing neurons in the ventral pallidum (Fig. 7Ha), which had dense terminals in the anterior portion of the LHb (Fig. 7Hb and L). Consistently, recording from LHb neurons in acute slices while photostimulating, we observed large (−8.8 ± 2.3 mV) GABAzine-senstive IPSPs in 6 of 13 recorded neurons (Fig. 7J and K), consistent with the solely inhibitory responses observed in data from our SOM-IRES-Cre::Ai32 transgenic line (Fig. 5K and L). Hence, taking this data altogether, we report two distinct source of extrinsic inhibitory input to the LHb, arising from PV-positive neurons in the MDT, and SOM-positive neurons in the VP.

## Discussion

In this study we sought to address the issue of inhibitory control within the LHb. We implemented the use of three markers known to represent distinct sub-populations of inhibitory neurons within the neocortex (Abs et al., 2018; Tasic et al., 2016; Tremblay et al., 2016), hippocampus (Klausberger and Somogyi, 2008) and striatum (Tepper et al., 2011). We provide evidence for three sources of inhibitory input to the LHb mediated by separate populations of PV-positive neurons within the LHb itself, and in the medial dorsal thalamus; and SOM-positive neurons in the ventral pallidum.

### PV neurons provide input to the LHb from a variety of sources

The subject of locally-targeting inhibitory neurons within the LHb has classically been a topic of debate (Brinschwitz et al., 2010; Weiss and Veh, 2011). In this work we show that at least two distinct populations of PV-positive neurons provide inhibitory input to the LHb; one locally-targeting and one located ventrally in the MDT. Strikingly, these neurons appear to selectively target neurons located within the lateral LHb; particularly the oval sub-nucleus (Figs. 7D and E). While the efferent targets of the LHb have long been known (Herkenham and Nauta, 1979), dissecting the output pathways with respect to distinct habenular sub-regions is a relatively novel idea (Quina et al., 2015). Indeed it is now known that topographical organisation of LHb outputs exist and it is believed that neurons in the lateral LHb specifically project to the rostromedial tegmental nucleus (RMTg) (Quina et al., 2015), the primary inhibitory modulator of the VTA (Jhou et al., 2009a, 2009b). Thus it is interesting to speculate whether these two distinct sources of inhibition converge specifically on RMTg-projecting neurons in the lateral LHb. If this were to be the case, theoretically these neurons would be well poised to reduce excitatory output to the RMTg and reduce the ‘reward aversion’ signalling from the LHb (Li et al., 2011; Matsumoto and Hikosaka, 2007). Further work employing *in vivo* optogenetic and chemogenetic manipulations in combination with behavioural testing could serve to answer these questions.

In addition to these two populations of PV-positive inhibitory neurons, our results also imply the existence of several physiologically diverse sub-populations of PV-positive neurons within the LHb (Fig. 4), which we assume to be projection neurons. These neurons possess clearly distinct physiological properties from the generic LHb neuron physiology (Chang and Kim, 2004; Kim and Chang, 2005; Weiss and Veh, 2011), and also appear to be roughly obey sub-nuclear boundaries (Andres et al., 1999). Thus we also speculate that these neurons may have differing projection targets and consequently differing functions. Further studies employing the use of Cre-dependent viral tracers are necessary to delineate the circuitry that these projection neurons comprise.

### Possible implications for inhibitory SOM-positive ventral pallidal neurons

We also report SOM-positive inhibitory projection neurons in the ventral pallidum which target the LHb (Fig. 7). Although inhibitory LHb-targeting ventral pallidal neurons have very recently been described (Faget et al., 2018; Stephenson-Jones et al., 2019), their function has remained largely elusive. Recent work suggests that LHb-projecting inhibitory and excitatory ventral pallidal neurons act oppositely to encode positive and negative motivational valence (Stephenson-Jones et al., 2019). By identifying that SOM acts as a specific molecular marker for these inhibitory neurons, we can thus facilitate further study of this pathway and more thoroughly investigate the role of the ventral pallidum to LHb pathway in controlling motivation.

### An absence of markers for studying the role of neurogliaform cells in the LHb

Ndnf has recently been shown to be expressed selectively by layer 1 neurogliaform cells in the neocortex (Abs et al., 2018; Tasic et al., 2018, 2016). In contrast we find no such evidence in the LHb (Fig. 4). Consistently, we observed no neuronal somata positive for NPY (Fig. S1). Therefore, our results do not support the notion of the existence of neurogliaform cells within the LHb (Wagner et al., 2016a; Weiss and Veh, 2011); or at least not those with similar molecular marker expression to those described in the neocortex (Abs et al., 2018; Tasic et al., 2018, 2016). However, these previous works were carried out in rat, hence we cannot exclude that this discrepancy is simply a species difference. We did observe GABA-immunoreactive Ndnf-positive LHb neurons. However, these only represent a small fraction of Ndnf-positive neurons (Fig. 2E). Hence, while it is therefore possible that neurogliaform cells are present within the mouse LHb, we conclude that neither Ndnf nor NPY can be used as a marker to study them.

### A need for sub-classifying LHb neurons based on physiological properties and molecular marker expression

Much invaluable information regarding the LHb has come from histological (Andres et al., 1999; Brinschwitz et al., 2010; Geisler et al., 2003) and transcriptomic studies (Aizawa et al., 2012; Wagner et al., 2016b, 2016a). These works have provided great insight into the organization of the LHb neuronal circuitry with reference to particular protein and gene expression patterns. However, these approaches do not permit these findings to be correlated with neuronal physiology at the single cell level. Indeed, recent reports have highlighted that individual LHb neurons which exhibit ‘bursting’ physiological phenotypes are associated with depression (Cui et al., 2018; Yang et al., 2018), and thus a greater understanding of LHb neuronal physiology, and how this physiology links to molecular protein expression is of great value.

Within this work, we have approached this problem by investigating the physiological properties of neurons expressing defined molecular markers. In the case of PV-positive LHb neurons alone, we show that these can be sub-classified into inhibitory neurons, and at least two distinct classes of non-inhibitory neurons (Fig. 4). We also show that SOM-positive LHb neurons represent a physiological sub-class with hybrid properties of LHb and MHb neurons (Fig. 4). What could be the functional importance of these neurons then? We now know that calcium-channel mediated bursting acts as a synchronization mechanism (Gutnick and Yarom, 1989; Wilcox et al., 1988), which in the case of the LHb acts to increase overall output to downstream structures and is thus critical in the pathogenesis of depression (Cui et al., 2018; Yang et al., 2018). Assuming that the hyperpolarisation-induced change in firing modality in these neurons is at least in part mediated by similar low-threshold channels, one could then speculate that this may also be some form of synchronization mechanism specific for these SOM-positive LHb neurons. It has also been proposed that this superior sub-nucleus of the LHb where these SOM-positive neurons are mostly confined (Fig. 2) may act as an area of interaction between the LHb and MHb (Wagner et al., 2016b). Assuming this to be true, perhaps then these neurons may act to synchronize activity between these two regions and increase overall habenular output, or perhaps they may have different downstream targets altogether from both the LHb and MHb.

As of yet, the specific functions of these PV-positive and SOM-positive neurons remain unclear. However, implementing the use of molecular markers as tools to study particular sub-populations of LHb neurons will most likely facilitate a greater understanding of habenular circuitry at the cellular level.

### The functional importance of inhibitory signalling within the LHb

It is well established that the LHb is hyperactive in MDD (Cui et al., 2018; Lecca et al., 2016; Li et al., 2011; Sartorius et al., 2010; Shabel et al., 2014; Tchenio et al., 2017; Yang et al., 2018), and that reducing LHb hyperexcitability by altering the balance of excitatory and inhibitory signalling has an antidepressant effect (Lecca et al., 2016; Li et al., 2011; Meye et al., 2016; Shabel et al., 2014; Wallace et al., 2017). A striking feature of many neurons innervating the LHb is that they co-release GABA and glutamate and hence much work has focussed on the role that these neurons play in modulating LHb excitability (Lazaridis et al., 2019; Meye et al., 2016; Root et al., 2014; Shabel et al., 2014; Wallace et al., 2017). Yet, both local (Zhang et al., 2018) and projection (Faget et al., 2018; Huang et al., 2019) GABAergic neurons are also known to innervate the LHb.

How these GABAergic neurons modulate the LHb in MDD remains to be fully established. Indeed, one could speculate that if excitatory inputs to the LHb are potentiated in MDD (Li et al., 2011), then GABAergic tone may be reduced. If this were to be the case, then one may also speculate that increasing GABAergic tone should also have an antidepressant effect. For this reason, we believe GABAergic innervation of the LHb to be an important topic of study. Interestingly, our results seem to point to a target selectivity of inhibitory PV-positive neurons at the sub-regional, possibly even single cellular level (Figs. 5I and 7D). We also show that these inhibitory neurons are capable of silencing the spontaneous activity of LHb neurons (Fig. 5J). Therefore, it is reasonable to assume that the role played by these inhibitory neurons in modulating LHb neuronal excitability is not negligible, and hence a greater understanding of this inhibitory control is an exciting concept.

## Conclusions

We have investigated the mechanisms of inhibitory control within the LHb and have defined three sources of inhibitory input from both locally-targeting PV-positive LHb neurons, and from those in the MDT; and from SOM-positive neurons in the VP. We also report multiple physiologically distinct populations of PV-positive and SOM-positive LHb neurons, and provide further evidence (Knowland et al., 2017; Wallace et al., 2017) for excitatory input to the LHb from PV-positive projection neurons (Fig. 5). These results, combined with the lack of specificity of Ndnf for neurogliaform cells within the LHb, also indicate that these markers represent broadly diverse populations of neurons on a region-to-region basis and therefore these populations must be validated in each case.

## Acknowledgements

We are grateful to Hongkui Zeng from the Allen Brain Institute, Seattle, for kindly sharing the Ai9 reporter mice with us, and we thank Csaba Földy, Brain Research Institute, University of Zurich, Switzerland, for generously providing us with NPY-hrGFP mice.

This work was supported by a Royal Society Research Grant, RG160549 (C.W.), a Wellcome Trust Seed Award, 205917/z/17/Z (C.W.), a Small Research Grant from Tenovus Scotland, Project S16/19 (C.W.), the DFG FOR2143 (P.W.), and BBSRC (BB/M00905X/1 to S.S.). J.F.W. holds a studentship funded by the Engineering and Physical Sciences Research Council (EPSRC).

We further thank the Wozny lab for helpful discussions. The authors have no financial conflicts of interest.

## Supplementary figures

**Supplementary Figure 1:**
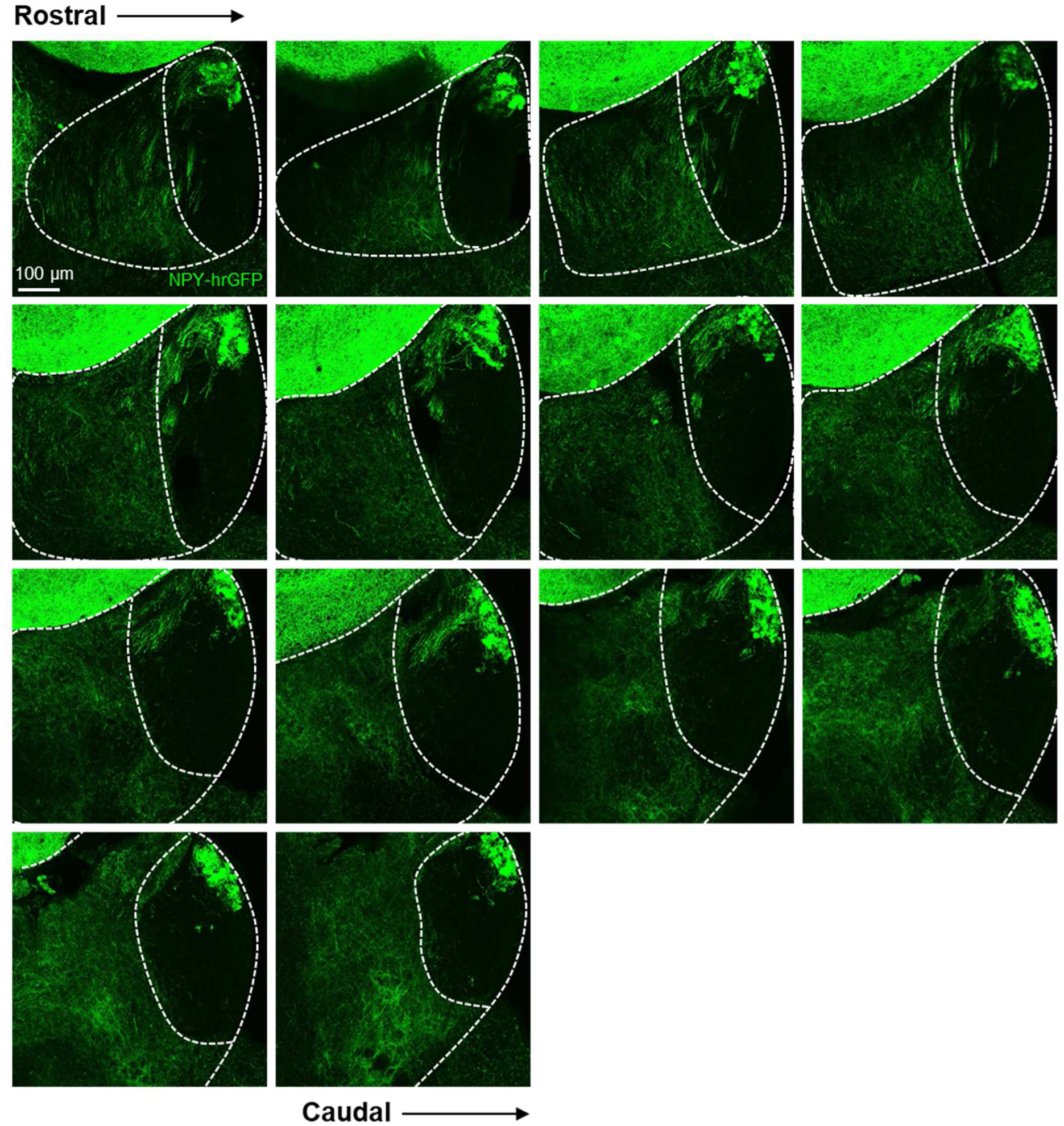
Absence of NPY-positive neuronal somata within in the LHb. Confocal micrographs of habenular sections from NPY-hrGFP mice (N = 2) depicting NPY-expression throughout the LHb in the rostral-caudal plane. Images are maximum intensity projections of 50 µm tissue. Note that no NPY-positive somata are located within the LHb.

**Supplementary Figure 2:**
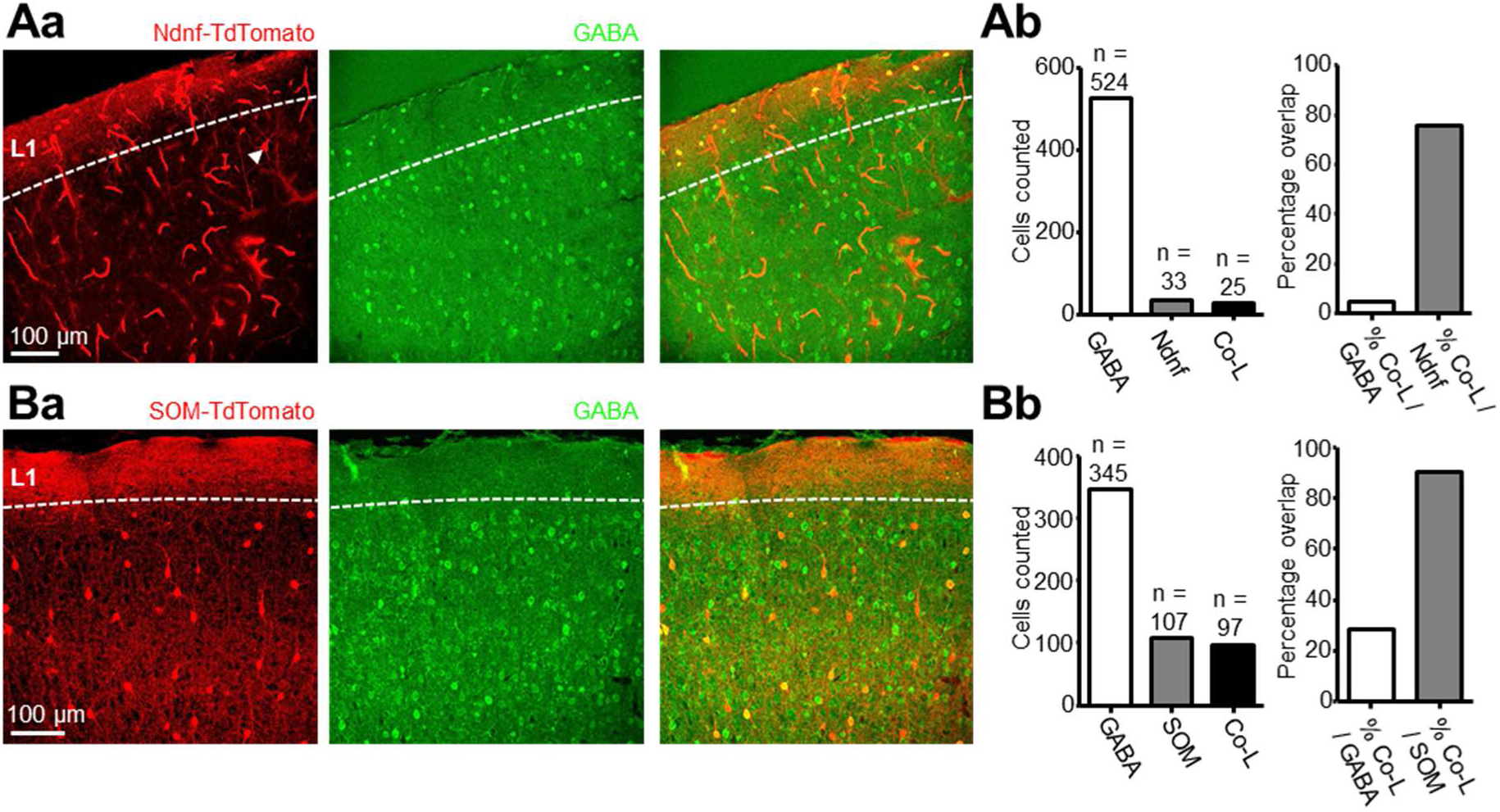
Ndnf and SOM are both expressed primarily by GABAergic neurons in the somatosensory cortex. **(Aa)** Confocal micrograph from Ndnf-IRES-Cre::Ai9 slices depicting Ndnf-positive neurons in L1 of the somatosensory cortex, but also an Ndnf-positive neuron in the deeper somatosensory cortex which was not immunoreactive for GABA. **(Ab)** Left: bar chart quantifying total number of GABA-immunoreactive, and Ndnf-positive neurons counted and the number which co-expressed both (N = 2 mice). Right: fraction of neurons expressing both markers as a percentage of GABA-immunoreactive neurons, and as a percentage of Ndnf-positive neurons. **(Ba)** As for (Aa), in slices from SOM-IRES-Cre::Ai9 mice (N = 2). **(Bb)** As for (Ab), in slices from SOM-IRES-Cre::Ai9 mice.

**Supplementary Figure 3:**
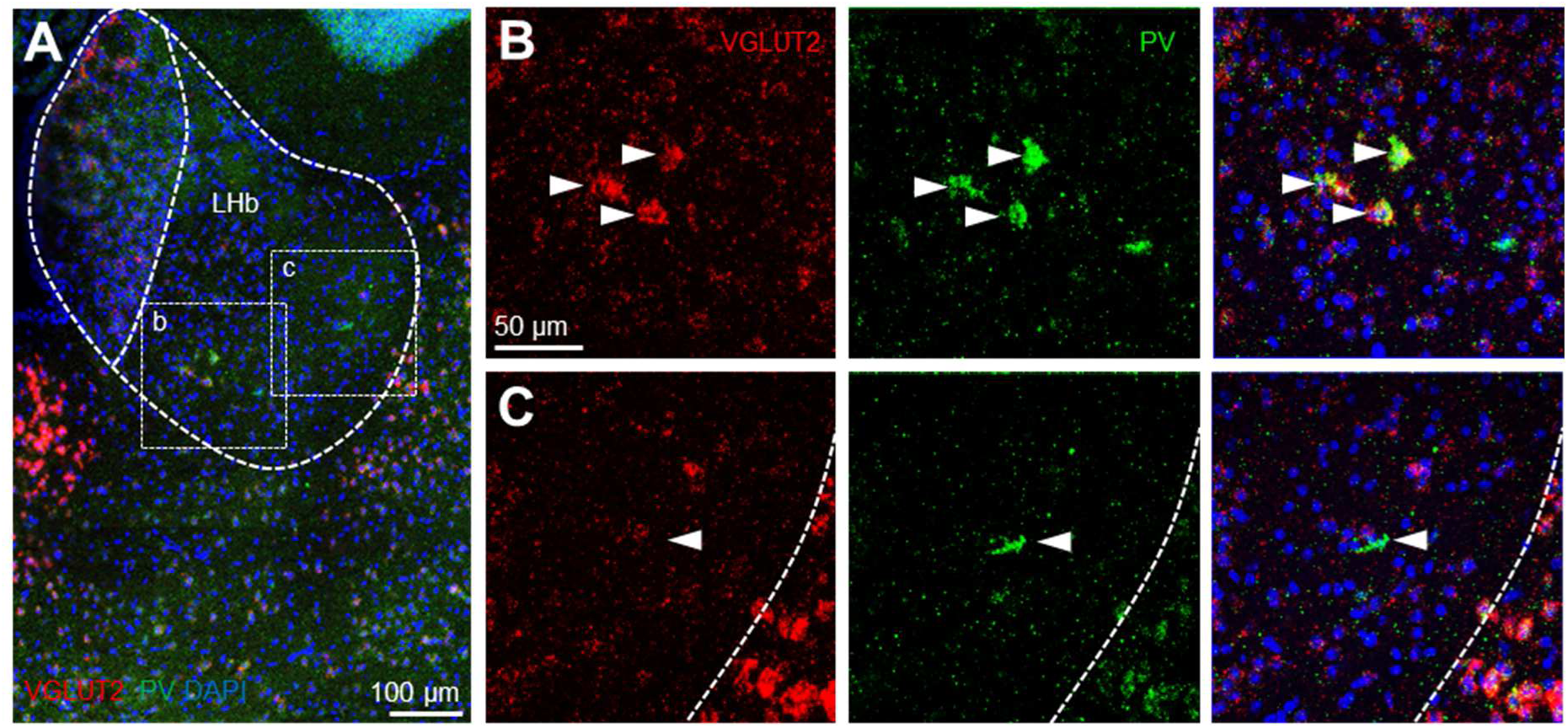
PV-positive neurons in the medial LHb are glutamatergic. **(A)** Overview image of the LHb from PV / VGlut2 double in situ hybridisation. **(B)** Zoom of boxed region in (A) depicting PV / Vglut2 double-positive neurons in the medial LHb. **(C)** Zoom of boxed region in (A) depicting PV-positive neurons which were negative for VGlut2.

**Supplementary Figure 4:**
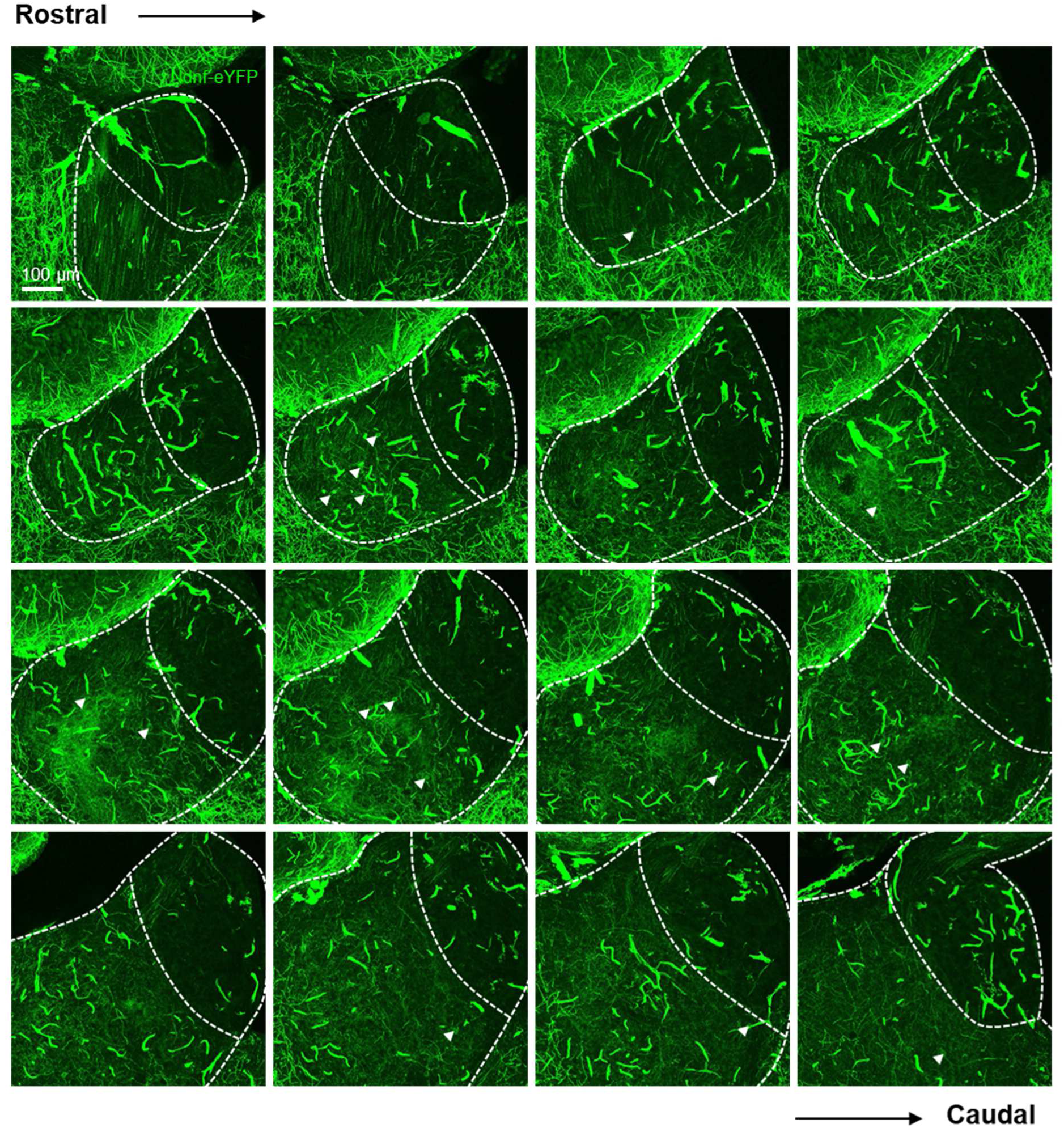
Localization of Ndnf-positive neuronal somata and processes throughout the LHb. Confocal micrographs of habenular sections from Ndnf-IRES-Cre::Ai32 mice (N = 2) depicting Ndnf-expression throughout the LHb in the rostral-caudal plane. Images are maximum intensity projections of the most superficial 30 µm of tissue from 60 µm slices. Arrowheads indicate Ndnf-positive neurons.

**Supplementary Figure 5:**
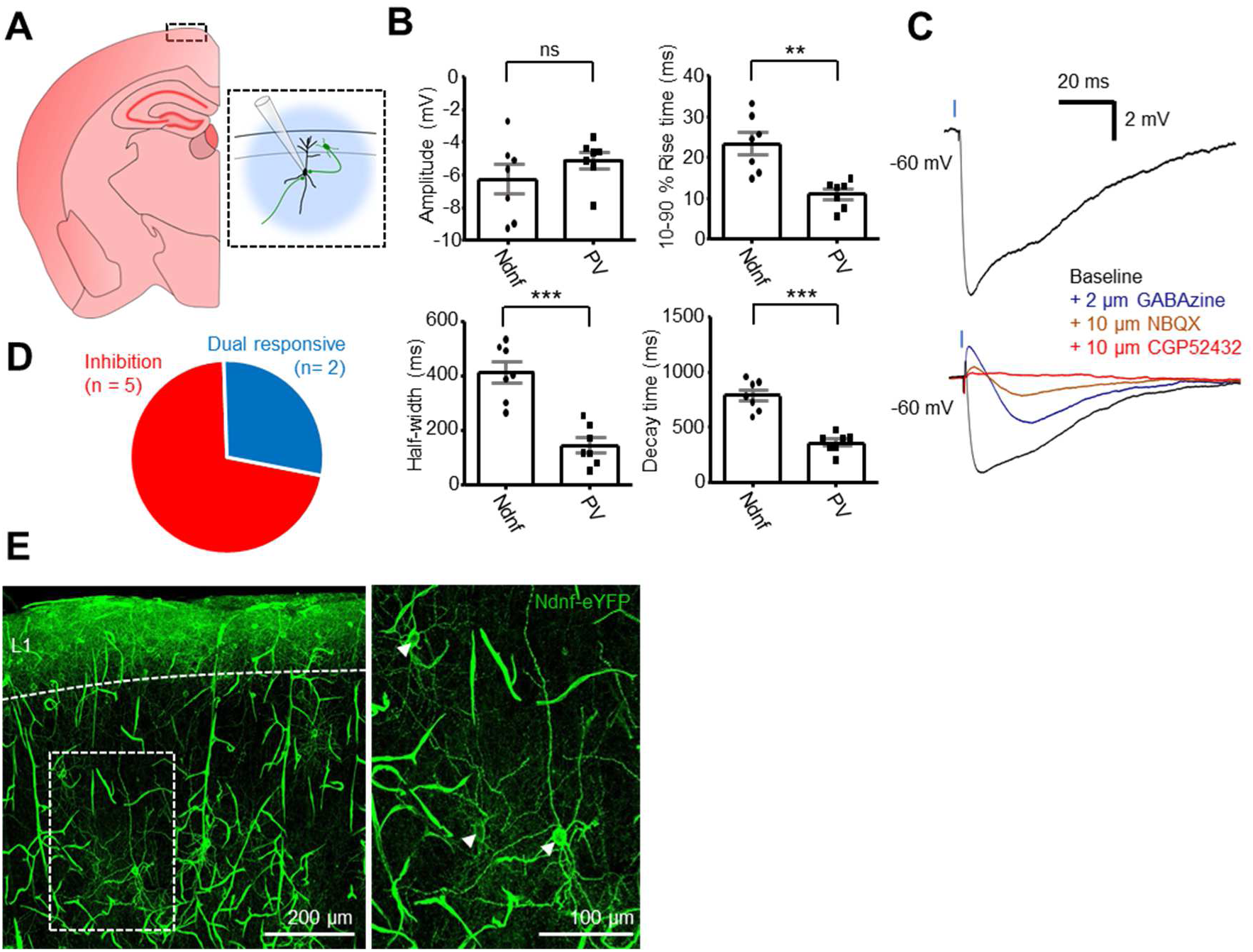
Ndnf is expressed selectively but not exclusively by L1 neurons in the somatosensory cortex. **(A)** Schematic of recording scheme. L2/3 pyramidal neurons were patched and blue light was used to stimulate both local and long-distance Ndnf-positive neurons. **(B)** Scatter plots comparing amplitude (−6.3 ± 0.9 mV vs −5.1 ± 0.5 mV; *p* = 0.29; two-tailed unpaired t-test) and kinetics (rise time 23.5 ± 2.7 ms vs 11.1 ± 1.3 ms; *p* = 0.001; half-width 414.6 ± 39.1 ms vs 146.0 ± 27.6 ms; *p* = 0.0001; decay 787.4 ± 51.9 ms vs 361.3 ± 32.8 ms; *p* < 0.0001; two-tailed unpaired t-test) of photostimulation-induced IPSPs in Ndnf-IRES-Cre::Ai32 slices vs PV-IRES-Cre::Ai32 slices. **(C)** Example traces from a neuron displaying an IPSP in response to photostimulation (top) and from one neuron in which an NBQX-sensitive EPSP was unmasked upon application of GABAzine (bottom). **(D)** Pie chart quantifying nature of postsynaptic potentials elicited in response to photostimulation. n = 7 neurons. **(E)** Left: Confocal micrograph depicting localization of Ndnf-positive neurons in the somatosensory cortex. Right: Zoom-in of left image showing Ndnf-positive neurons not located within neocortical L1 (arrowheads).

**Supplementary Figure 6:**
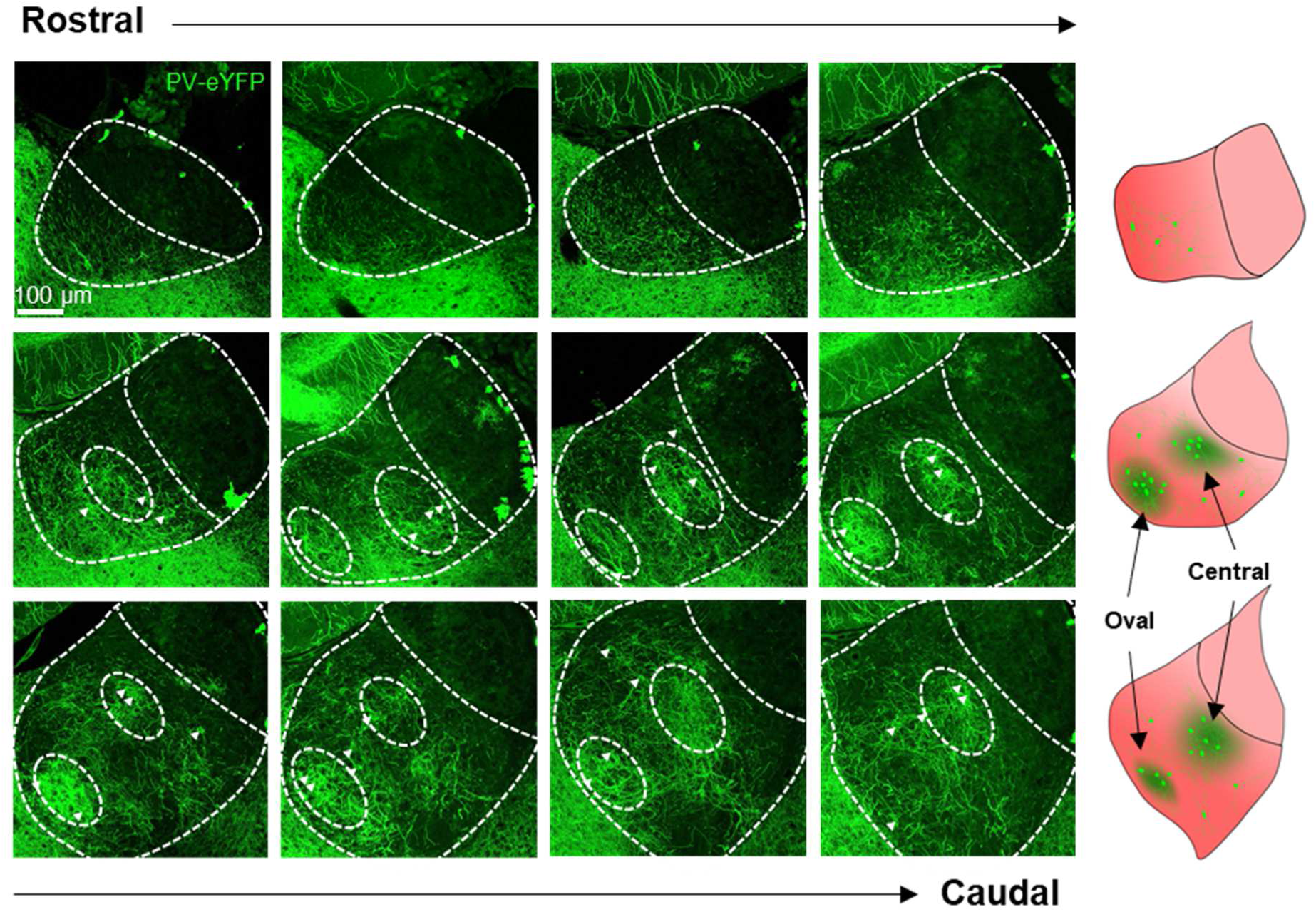
Localization of PV-positive neuronal somata and processes throughout the LHb. Left: Confocal micrographs of 30 µm thick maximum intensity projections of habenular sections from PV-IRES-Cre::Ai32 mice (N = 2) depicting localization of PV-positive neuronal somata and neuronal processes throughout the LHb in the rostral-caudal plane. 60 µm thick spacing between images. Arrowheads indicate PV-positive neurons. Right: graphical illustrations of habenular sections indicating the oval and central lateral habenular sub-regions, where PV-expression was most prominent. Images each represent one third of the habenula in the rostral-caudal plane.

**Supplementary figure 7:**
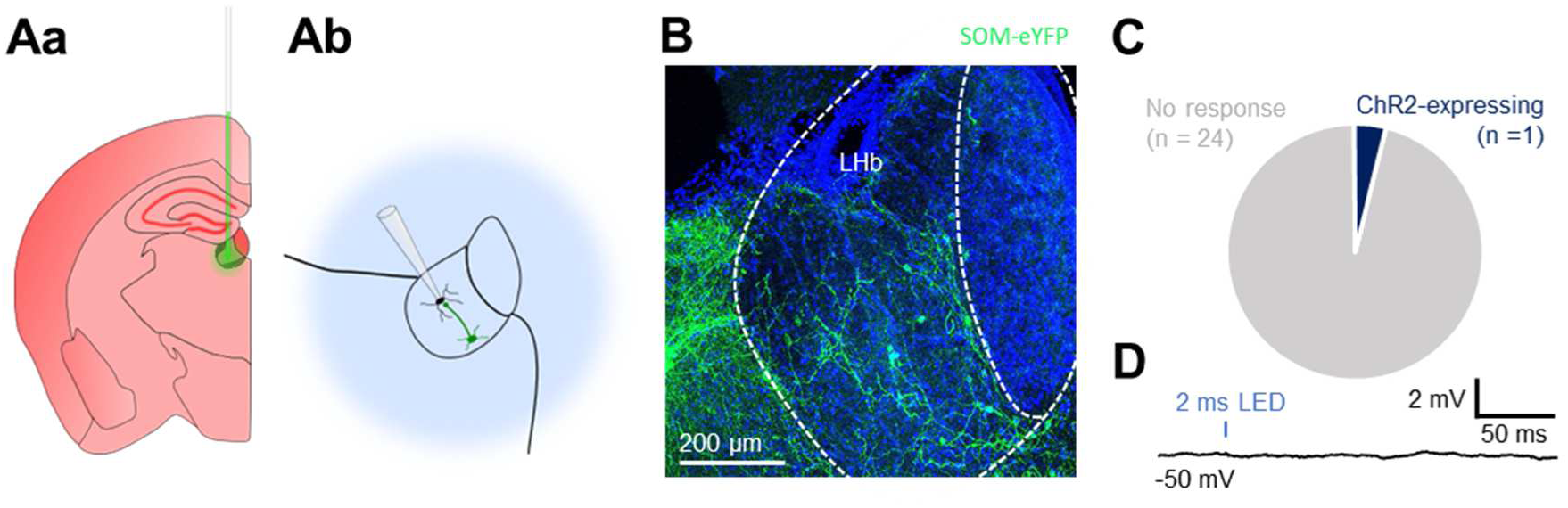
SOM-positive LHb neurons do not mediate local inhibition. **(Aa)** Schematic illustrating stereotaxic injection protocol of AAV9 into the LHb of SOM-IRES-Cre mice (N = 4). **(Ab)** Schematic illustrating electrophysiology recording protocol for LHb neurons following stereotaxic viral injection. Transduced SOM-positive neurons are photostimulated while recording from nearby LHb neurons. **(B)** Confocal micrograph depicting virally-transduced SOM-positive neurons within the LHb. **(C)** Pie chart quantifying fraction of neurons responsive to photostimulation. **(D)** Representative example trace from a neuron which showed no response to photostimulation. Trace is an average of multiple individual sweeps.

**Supplementary figure 8:**
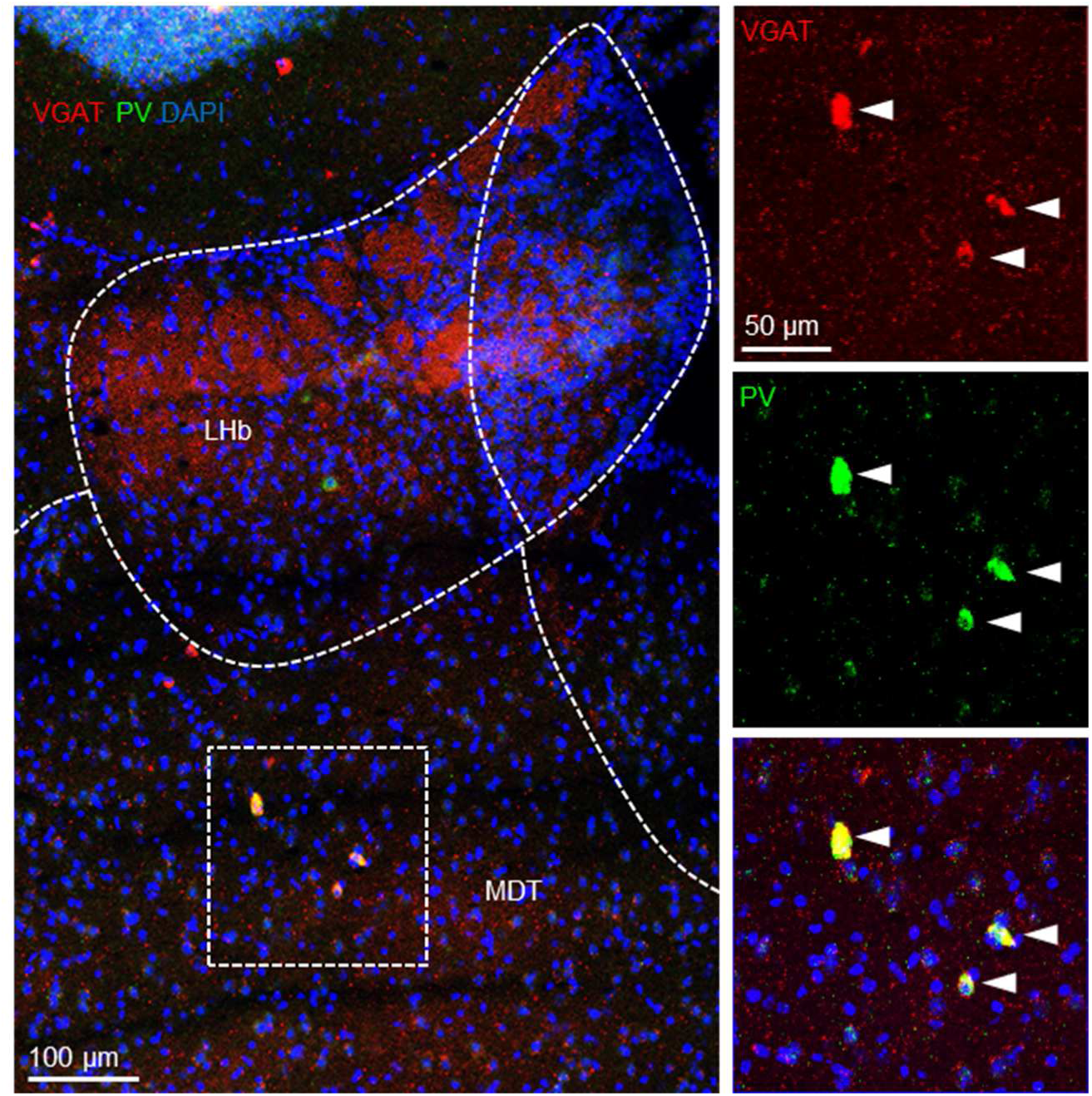
PV-positive neurons in the MDT are GABAergic. In situ hybridization depicting VGAT / PV double positive neurons in the MDT.

